# Classical macrophage polarisation is limited by human β-defensin-3 via an autocrine IL-4 dependent process

**DOI:** 10.1101/2021.05.06.442606

**Authors:** Maria E. Candela, David J.P. Allsop, Roderick N. Carter, Fiona Semple, Fiona Kilanowski, Sheila Webb, David Taggart, Henry J.W Mullan, Brian J. McHugh, David H. Dockrell, Donald J. Davidson, Judith E. Allen, Stephen J. Jenkins, Nicholas M. Morton, Julia R. Dorin

## Abstract

Human β-defensin 3 (HBD3), is an anti-microbial host-defence peptide, that can rapidly enter macrophages to modulate TLR4 responses to lipopolysaccharide. However, the molecular mechanisms by which HBD3 exerts this anti-inflammatory influence remain unclear. Here, we show mice deleted for the orthologue of HBD3 have an increased acute lipopolysaccharide response *in vivo*. Furthermore, we found that HBD3 limited the response of macrophages to classical activation, and contemporaneously drove expression of IL-4. An increase in markers of alternative activation, and a change in metabolic flux was also observed. Consistent with these results, HBD3 enhanced the IL-4 mediated polarisation of naïve macrophages. Finally, we demonstrate that the ability of HBD3 to limit macrophage classical activation requires IL-4Rα. These data reveal a previously unrecognised role for HBD3 in influencing the polarisation state of macrophages to enable a state conducive for repair and resolution.

**SYNOPSIS:** 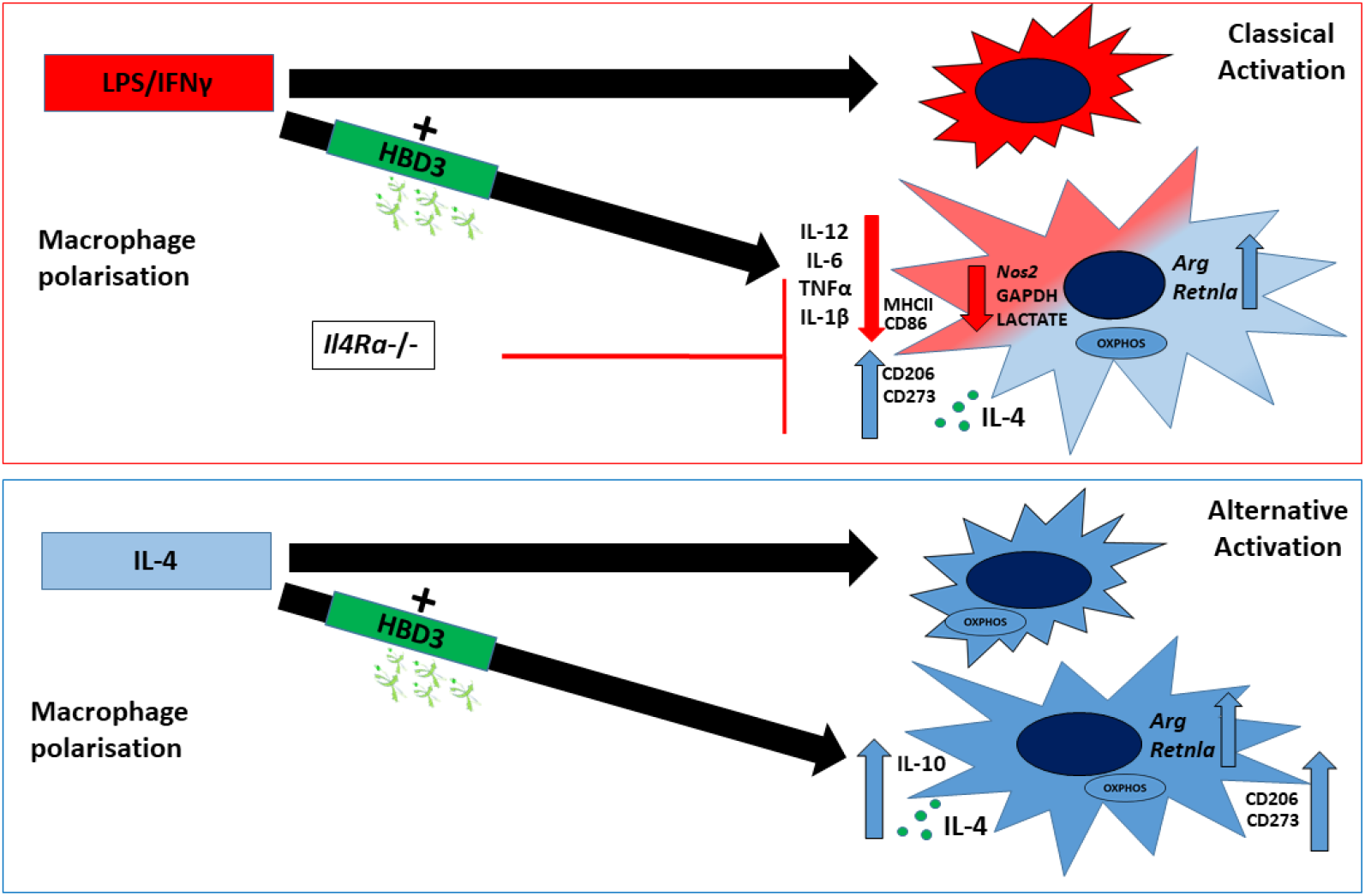

The anti-microbial host-defence peptide, Human β-defensin 3 (HBD3), is shown here to modulate the inflammatory response to classical activation by promoting alternative activation through IL-4Rα, to enable a state conducive for repair and resolution.

- Knockout mice for the orthologous gene for HBD3, demonstrate increased acute lipopolysaccharide inflammatory response.
- HBD3 limited the classical activation of macrophages polarised with LPS/IFNγ and drove expression of IL-4. Cells also displayed increase in alternative activation markers and promotion of oxidative phosphorylation.
- HBD3 enhanced the IL-4-mediated activation of naïve macrophages.
- The ability of HBD3 to limit macrophage classical activation and contemporaneously promote alternative activation required IL-4Rα.

## Introduction

The β-defensins are a multigene family encoding antimicrobial, cationic, amphipathic peptides. They have a conserved structure stabilized by a canonical six cysteine motif and specific disulphide connectivities (Bauer *et al*, 2001; Wu *et al*, 2003). Their expression at surface epithelia is rapidly induced in response to inflammatory signals, and several are expressed in response to vitamin D, lipopolysaccharide (LPS) and/or interferon γ (IFNγ) (Duits *et al*, 2002; Edfeldt *et al*, 2010). HBD3, encoded by *DEFB103*, is one of many β-defensins present in humans. Although mice have species-specific β-defensin clades, *Defb14* is the clear orthologue of *DEFB103*, and the peptides are 64% identical at the amino acid level (Schutte *et al*, 2002). Defensins have been termed alarmins, as molecules that can activate and mobilise immune cells in response to danger (Yang *et al*, 2013). Indeed, both HBD3 and DEFB14 are chemo-attractive for macrophages through CCR2. In addition, when HBD3 is added to macrophages, it is detected inside the cell within ten minutes (Rohrl *et al*, 2010; Semple *et al*, 2011). It has been shown that HBD3 increases the endosomal TLR9 response to pathogen or self-DNA in plasmacytoid dendritic cells (Lande *et al*, 2015; Lee *et al*, 2019; McGlasson *et al*, 2017; Rohrl *et al*., 2010; Tewary *et al*, 2013), and the macrophage response to high molecular weight Poly(I:C) via the cytoplasmic MDA5 (Semple *et al*, 2015). However, HBD3 has a dichotomous behaviour, and in addition to amplifying the response to some pattern recognition receptor ligands, the cytokine response to other molecular patterns can be reduced (Shelley *et al*, 2020).

We have previously shown that HBD3 reduces the TLR4-dependent transcriptional signature and cytokine response to LPS *in vitro* and *in vivo* (Semple *et al*, 2010). The causative mechanism is unknown, but it is neither due to LPS neutralisation, nor the binding and blocking of TLR4 by the peptide. The evidence for this is twofold, firstly LPS stimulation of TLR4 and assembly of the MyD88 signalling hub (MyDDosome) is known to happen within minutes of TLR4 stimulation (Latty *et al*, 2018), and the suppressive effect of the defensin on cytokine production is still evident even when peptide is added up to 60 minutes after LPS exposure (Semple *et al*., 2010). Secondly, NF-kB signalling induced by exogenous MyD88 expression and therefore independent of TLR4, is also reduced by HBD3 expression (Semple *et al*., 2011).

Here, we show that the serum cytokine response to *E*.*coli* LPS, is increased in mice with a targeted disruption of *Defb14*. In addition, we show that in the presence of HBD3 the degree of polarisation to classical activation by LPS and IFNγ is reduced, and there is an increase in alternative activation. These changes are coincident with increased IL-4 expression. Independently, HBD3 used in combination with IL-4 augments macrophage polarisation to alternative activation. A change in metabolic flux to oxidative phosphorylation is also observed when HBD3 is present in the classical activation polarisation, together with a reduction in expression of genes involved in aerobic glycolysis. Finally we show, that the effect of HBD3 on macrophage polarisation with LPS/IFNγ is dependent on IL-4Rα. This work extends the functional repertoire of β-defensins and provides mechanistic insights, with relevance for infection resolution and a return to homeostasis.

## Results

### HBD3 limits the pro-inflammatory M(LPS+IFNγ) polarisation phenotype of macrophages in vitro and in vivo

We have previously demonstrated that *in vitro*, DEFB14 and HBD3 peptides suppress lipopolysaccharide (LPS) signalling in primary macrophages (Semple *et al*., 2011). To determine if the effect of suppressing the pro-inflammatory cytokine response to LPS is relevant *in vivo*, we disrupted the *Defb14* gene using embryonic stem cell gene targeting (**Supplementary Fig.1**). We show here that, 90 minutes following LPS intraperitoneal injection, *Defb14* ^*tm1a*^ homozygotes had significantly greater increase in levels of IL-6 in their sera, compared to wild type animals (**Fig. 1a**). Thus indicating that in the absence of *Defb14* the acute response to LPS was increased *in vivo*. This emphasises the physiological relevance of β-defensin suppression of pro-inflammatory cytokines. We then exposed wild type, naïve bone marrow derived macrophages (BMDM) to the more potent classical activation stimulus of LPS and IFNγ (termed here M(LPS/IFNγ (Murray *et al*, 2014)), and followed this, with varying concentrations of HBD3 peptide. Macrophages polarised in the presence of (LPS/IFNγ) and HBD3 after 15 minutes (termed here M(LPS/IFNγ+HBD3), secreted significantly less TNF-α after 18 hours (**Fig. 1b)**. The suppressive effect was not significant at peptide levels less than 2.5μg/ml, and was most potent at 10μg/ml of peptide. This effect of HBD3 was reduced at 20μg/ml and lost at 30μg/ml (**Supplementary Fig. 2a**), which may reflect the cytotoxic effect of HBD3 on mammalian cells at higher concentrations (Leelakanok *et al*, 2015). The effect was structure dependent as addition of a linear peptide with an equal charge but lacking a disulphide-stabilised structure, did not reduce the TNF-α cytokine response. For all future experiments we used 5µg/ml (1µM) HBD3.

**Figure 1:**
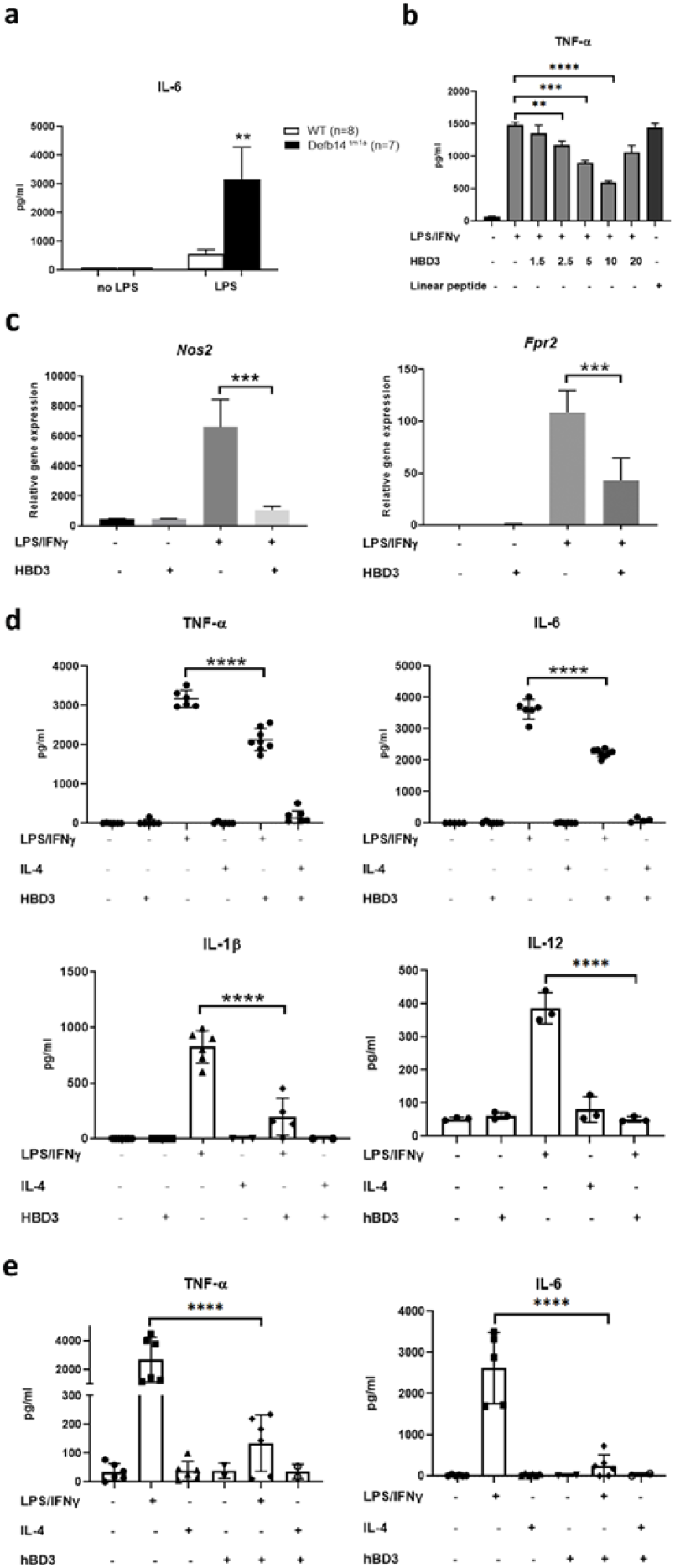
Defensin HBD3 limits the classical inflammatory polarisation phenotype of macrophages in human and mouse in vitro and in vivo. BMDM polarised as shown with or without (5 ug/ml) HBD3 after 15-30 minutes (except B where concentrations are shown) for 18 hours. **a:** IL-6 levels in serum of Defb14-/-and wild type mice 90 minutes after LPS injection. ** p<0.01. **b:** TNF-α expression in supernatant of BMDMs Mθ and M(LPS+IFNγ) treated with different HBD3 concentration (or 5 ug/ml of Linear Peptide) for 18 hours, measured by ELISA. Sample reference M(LPS+IFNγ), **** p<0.0001, *** p<0.001, ** p<0.01. **c:** *iNos* and *Fpr2* gene expression were measured by real time RT-PCR. Results are normalized to mRNA expression in naïve macrophages. *** p<0.001. **d:** TNF-α, IL-6, IL-1β and IL-12 cytokine expression measured by ELISA. **** p<0.0001. **e:** TNF-α and IL-6 cytokine expression measured by ELISA in human macrophages, polarized as shown, with or without (5 ug/ml) HBD3. **** p<0.0001. Comparison were done with one-way ANOVA test.

M(LPS/IFNγ+HBD3) from mouse BM showed significantly reduced expression compared to M(LPS/IFNγ), of two key classical activation genes-*Nos2* (inducible nitric oxide synthase gene) and *Fpr2* (Formyl Peptide Receptor 2 gene) (**Fig. 1c**). The cytokine levels of TNF-α, IL-6 and IL-12 were also less in M(LPS/IFNγ) with HBD3, compared to without (**Fig. 1d**). As well as measuring less secreted IL-1β, we also detected less *Caspase 1 (Casp1)* gene expression in M(LPS/IFNγ+HBD3) implying inflammasome engagement was reduced in the presence of HBD3, consistent with a reduction in glycolysis (**Supplementary Fig. 2b**) and IL-4 involvement (Czimmerer *et al*, 2018; Moon *et al*, 2015). We also found peripheral blood monocyte derived macrophages (PBMDM) from healthy volunteer donors, stimulated with LPS/IFNγ, showed a greater than ten-fold reduction of TNF-α and IL-6 when also in the presence of HBD3 (**Fig. 1e**). HBD3 alone, and/or IL-4 had no effect on the secretion of pro-inflammatory cytokines (**Fig 1d-e**). When culturing the mouse BMDM, we noticed that the population of M(LPS/IFNγ+HBD3) had cells with the dendritic morphology characteristic of M(LPS/IFNγ) but also some more rounded cells reminiscent of IL-4 alternatively activated macrophages(Ploeger *et al*, 2013) (termed here M(IL-4)(Murray *et al*., 2014)) (**Supplementary Fig. 2c**). To investigate this further, we examined markers for both classical and alternative activation states in the M(LPS/IFNγ+HBD3) cell population.

### HBD3 increases alternative activation markers in M(LPS+IFNγ) and augments M(IL-4) polarisation

The co-stimulatory marker CD86, and MHC II are expressed at high levels in M(LPS/IFNγ) macrophages (Van den Bossche *et al*, 2016), and we found these levels were significantly lower in cells polarised with LPS+IFNγ followed by HBD3, with the number of double positive cells reducing by half (**Fig. 2a** and **Supplementary Fig 3a**). In contrast, the levels of markers associated with M(IL-4), such as CD273 (encoding programmed death ligand 2 (PDL2)) and CD206 (Mannose receptor) (Gundra *et al*, 2014; Huber *et al*, 2010) were increased significantly (91% of cells stained double positive for both) (**Fig. 2b** and **Supplementary Fig 3a**). IL-10 is known to be important in the TLR4 response and is induced after 6 hours in pro-inflammatory macrophages to limit the pro-inflammatory activation of cells (Ip *et al*, 2017). At 18 hours, we found M(LPS/IFNγ) expressed a high level of IL-10, and M(LPS/IFNγ+HBD3) also had a strong IL-10 response, but this was reduced in comparison to M(LPS/IFNγ) (**Fig 2c**). Conversely IL-10 levels were significantly increased in M(IL-4+HBD3) compared to M(IL-4) (**Fig 2c**). We also observed an increase in gene expression of *Il4* in BMDM cells polarised with (LPS/IFNγ+HBD3) compared to M(LPS/IFNγ) (**Fig 2d**). We found this was also true when we used the mouse macrophage cell line RAW 264.7, and cytokine levels of IL-4 were increased in supernatant from human peripheral blood monocyte derived macrophages treated with LPS/IFNγ+HBD3 (**Fig 2d**). These results demonstrate that when macrophages are polarised with LPS/IFNγ and HBD3, classical activation markers are reduced, but IL-4 expression and alternative activation markers (consistent with those present on M(IL-4)), are increased

**Figure 2:**
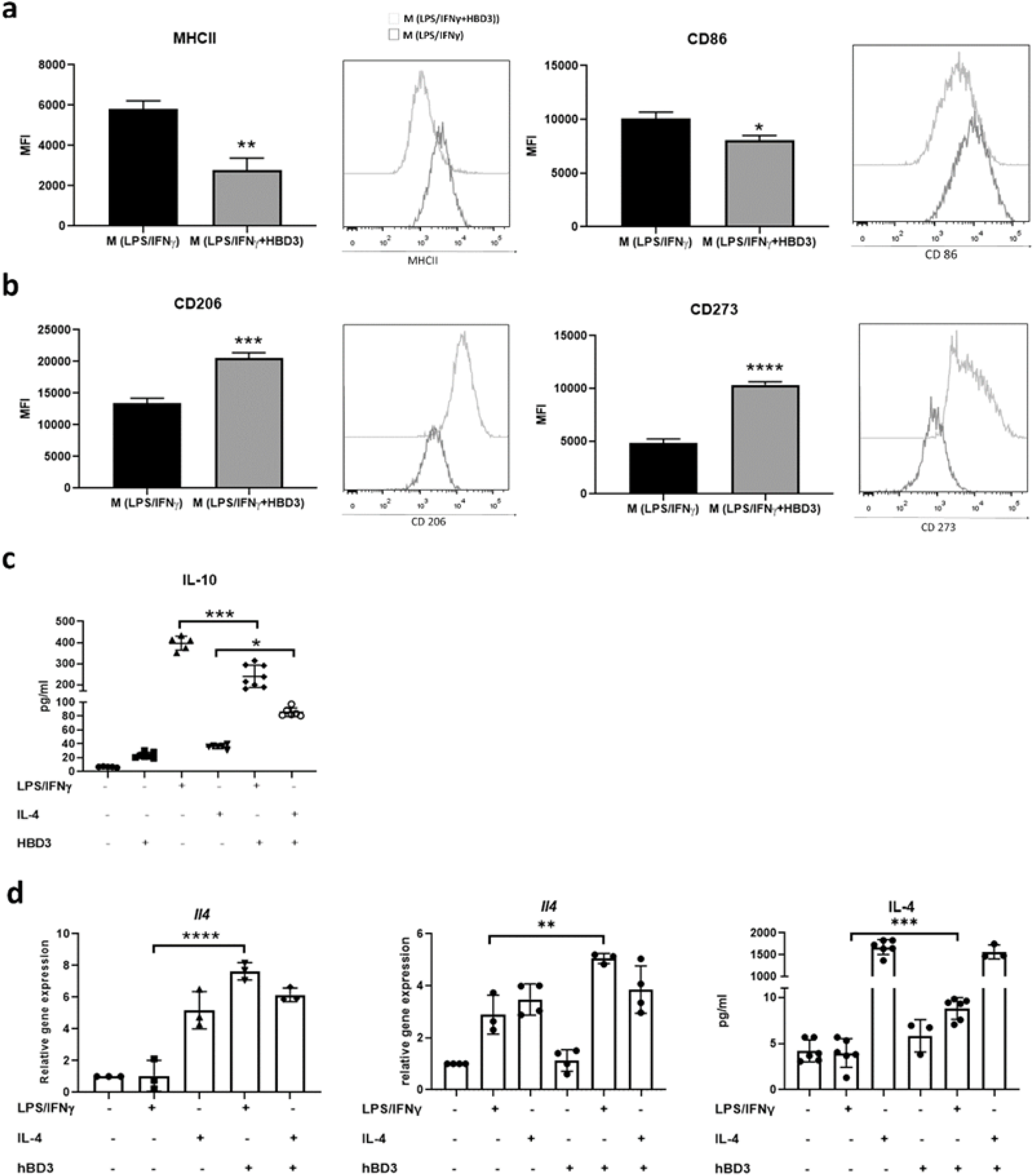
Decreased pro-inflammatory and increased alternative activation markers in M (LPS+IFNγ) in presence of HBD3. BMDM (samples from Fig. 1c-d), RAW 264.7 cells and human PBMDM (samples from fig. 1e) were polarised for 18 hours with LPS+IFNγ or IL-4 followed after 15-30 minutes +/-HBD3. **a-b:** Representative histograms of geometric mean fluorescence intensity (MFI) of MHCII, CD86, CD206 and CD273 surface markers analysed by flow cytometry. See **S3a for dot plot scattergrams.** **c:** IL-10 cytokine expression measured by ELISA. **d:** *Il4* gene expression measured by real time where the results are normalized to mRNA expression of naïve macrophages, in BMDM and RAW cells (left to right) and IL-4 cytokine in supernatant of human macrophages (far right). Comparison were done with one-way ANOVA test, **** p<0.0001, *** p<0.001, ** p<0.01, * p<0.05.

Given the ability of HBD3 to promote *Il4* expression in M(LPS/IFNγ), we speculated that HBD3 could enhance IL-4 alternative activation of macrophages. *Arginase 1* (*Arg1*) and *Resistin like molecule alpha* (*Retnla*), were both expressed at higher levels in M(IL-4+HBD3) compared to M(IL-4), when 20ng of IL-4 was used for polarisation (**Fig. 3a**). Co-culture with HBD3 did not affect the high levels of macrophage expression of CD206 or CD273 stimulated by 20ng/ml of IL-4 (**Fig. 3b**), however, this level of IL-4 may induce maximal activation, as has been shown in CD4+ Th2 cells (Perona-Wright *et al*, 2010). When we reduced the level of IL-4 stimulation to 5ng/ml or 10ng/ml, HBD3 induced a strong and significant augmentation of the gMFI of CD206 and CD273 and increase in the number of double positive cells (**Fig 3b** and **Supplementary Fig. 3b**). In addition, the fact that HBD3 augmented IL-4 polarisation of naïve BMDM to alternative activation, confirms again that the suppression of classical activation and increase in alternative activation markers LPS in response to LPS/IFNγ, is not mediated through LPS neutralisation. We then investigated whether the presence of HBD3 influences the cellular metabolic programming of macrophages.

**Figure 3:**
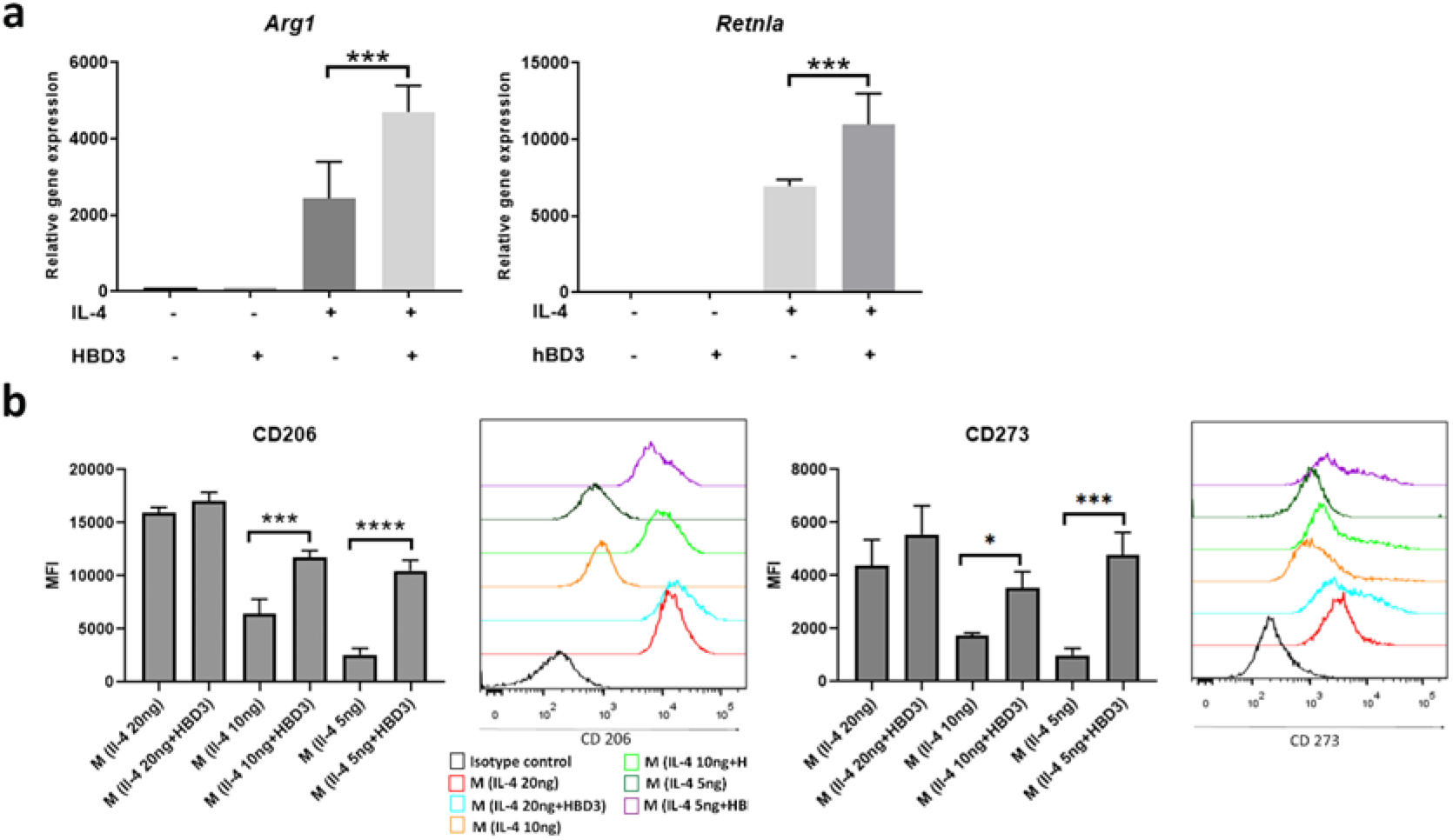
The presence of HBD3 augments IL-4 polarisation. **a:** *Arg1, Retnla* gene expression were measured by real time in cells polarised with and without IL-4 (20ng/ml) and with or without HBD3 as shown. Results are normalized to mRNA expression of naïve macrophages. *** p<0.001. **b:** Representative histograms of geometric mean fluorescence intensity (MFI) of CD206 and CD273 surface markers expression analysed by flow cytometry on cells stimulated for 18hrs with IL-4 at 5 or 10ng/ml. **** p<0.0001, *** p<0.001, * p<0.05. See **S3b for dot plot scattergrams**. Comparison were done with one-way ANOVA. **** p<0.0001, *** p<0.001, ** p<0.01, * p<0.05.

### M(LPS+IFNγ) show promotion of OXPHOS in the presence of HBD3

Pro-inflammatory macrophages characteristically utilise glycolysis for rapid energy production, while oxidative phosphorylation (OXPHOS) predominates in alternatively activated macrophages at 18-24 hours (O’Neill & Pearce, 2016). It is recognised that in response to inflammatory activators, such as LPS/IFNγ, glycolysis generates ATP, pyruvate and lactate from glucose, and the mitochondrial matrix associated tricarboxylic acid (TCA) or Krebs cycle is modified with specific breaks that allow build-up of certain intermediates and an elevation of mitochondrial membrane potential (Viola *et al*, 2019). Thus, mitochondria are repurposed from ATP production by OXPHOS, to succinate-dependent ROS generation (Mills *et al*, 2016). In contrast, alternatively activated macrophages such as M(IL-4), require the intact TCA cycle for OXPHOS, and promotion of the non-oxidative pentose phosphate pathway to satisfy the requirements for nucleotide synthesis, and N-glycosylation for alternative activation cell surface markers, such as CD206 (Jha *et al*, 2015; Wang *et al*, 2018). In agreement with this, we found naïve BMDM macrophages and those stimulated with IL-4, exhibited a high basal rate of oxygen consumption (OCR), indicative of OXPHOS, whereas LPS/IFNγ stimulated BMDM, displayed much lower OCR indicative of reduced OXPHOS **(Fig 4 & Supplementary Fig. 4**). Responses to mitochondrial stress induced by various inhibitors of the electron transport chain, were determined in BMDM, under the different polarisation conditions with or without HBD3 (**Fig. 4 and supplementary Fig 4**.). As expected, M(LPS/IFNγ) had reduced ATP respiration and maximal respiratory capacity levels compared to M(IL-4), indicating a lack of OXPHOS. The basal, ATP and maximal respiration measurements were all significantly higher in M(LPS/IFNγ+HBD3) than those in M(LPS/IFNγ). HBD3 alone did not influence basal, ATP or maximal respiration of M(IL-4) and M(θ). The lack of HBD3’s effect on M(θ) metabolism, mirrors the lack of effect HBD3 has on cytokine secretion or gene expression (**Figs 1 and 2**).

**Figure 4:**
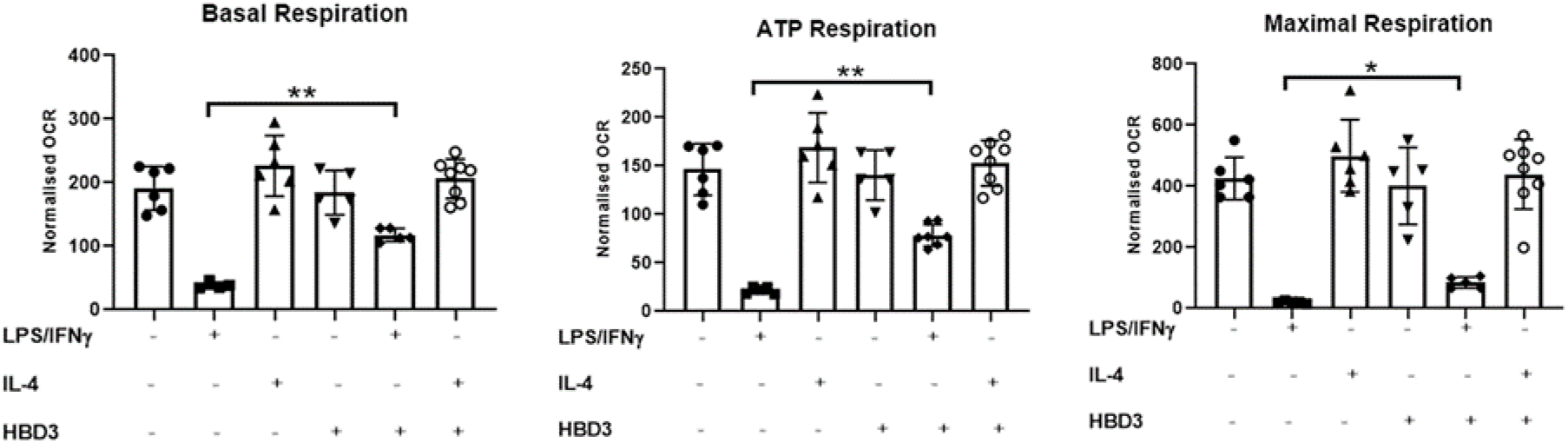
Mitochondrial respiration in M(LPS/IFNγ+HBD3). BMDM were polarised as labelled, with or without HBD3 for 18 hours. Representative experiment of cellular respiration measured under mitochondrial stress condition. Calculated basal respiration, ATP production and maximal respiration measured under mitochondrial stress condition. ** p<0.01, * p<0.05. Comparison were done with one-way ANOVA test. See **supplementary figure 4** for representative experimental graph.

Consistent with an increase in OXPHOS in M(LPS/IFNγ+HBD3), we also detected significant reduction in gene expression of the glucose transporter *Glut1*, the rate controlling enzymes in glycolysis *Glyceraldehyde 3-phosphate dehydrogenase* (*Gapdh*), and *Hexokinase 2* (*Hk2*), compared to M(LPS/IFNγ) (**Fig. 5a**). This partial change in cellular metabolism compared to that observed in M(LPS/IFNγ) reflected the partial reduction in pro-inflammatory cytokines that we observed (**Fig. 1**). Exogenous expression of *Glut1* in M(LPS) has been shown to increase glycolysis and pro-inflammatory cytokine production of TNF-α (Freemerman *et al*, 2014). We wished to check that the suppressive effect of HBD3 on inflammatory macrophages was not due to cell death, and staining cells with Annexin V and Propidium iodide (PI), revealed that M(LPS/IFNγ), M(IL4) and M(θ) showed equivalent numbers of live cells when polarised with or without HBD3 (**Fig. 5b**). Measurement of MTT (3-(4,5-Dimethylthiazol 2-yl)-2,5-diphenyltetrazolium bromide) absorbance however, showed decreased activity in M(LPS/IFNγ), compared to M(LPS/IFNγ+HBD3) (**Fig. 5b**). The MTT absorbance values in M(IL-4), M(IL-4+HBD3) and M(LPS/IFNγ+HBD3) were equivalent. This difference between M(LPS/IFNγ) and M(IL-4) has previously been described and postulated to reflect a loss of succinate dehydrogenase due to disruption of the TCA cycle in the former (Van den Bossche *et al*., 2016). Macrophages polarised with (LPS/IFNγ)+HBD3 did not show any cell death or decrease in MTT assay activity, suggesting retention of an intact TCA cycle, and consistent with the reduced secretion of IL-1β in M(LPS+IFNγ+HBD3) compared to M(LPS+IFNγ) (**Fig 1d**). We then looked at the activity of glyceraldehyde-3-phosphate dehydrogenase (GAPDH), a key enzyme in aerobic glycolysis, that catalyses the conversion of glyceraldehyde 3-phosphate to D-glycerate 1,3-bisphosphate in the enzymatic conversion pathway from glucose to pyruvate. Enzymatic activity of GAPDH in M(LPS/IFNγ+HBD3) was significantly lower than in M(LPS/IFNγ), and equivalent in Mθ, where glycolysis is not predominant. The low level of GAPDH activity was the same as the level present when the GAPDH inhibitor dimethyl fumarate (DMF) was used in the LPS/IFNγ polarisation (M(LPS/IFNγ+DMF) (**Fig 5c**). M(LPS/IFNγ+HBD3) had similarly reduced levels of GAPDH activity as M(LPS/IFNγ+DMF), also consistent with HBD3 altering glycolytic metabolic flux. We also found a significant reduction in GAPDH activity when M(LPS/IFNγ) were treated with a different antimicrobial peptide, cathelicidin (also called LL-37). In addition to reduced GAPDH activity, lactate levels were decreased in M(LPS/IFNγ+HBD3) and M(LPS/IFNγ+LL-37) compared to M(LPS/IFNγ), and were equivalent to the level in M(θ) and M(Il-4) (**Fig 5c**). Intracellular lactate levels rose in M(LPS/IFNγ) at this time point of 18 hours compared to naïve M(θ) cells, a feature expected from an increase in aerobic glycolysis, as the pyruvate generated can enter mitochondria, or be reduced in the cytoplasm to lactate (Ryan *et al*, 2019; Viola *et al*., 2019). The reduction of lactate levels in M(LPS/IFNγ+HBD3) was consistent with reduced glycolysis compared to M(LPS/IFNγ), and coincident with the promotion of OXPHOS, revealed by the mitochondrial stress experiments (**Fig. 4**). Taken together these results indicate that HBD3 can alter the metabolic programming of pro-inflammatory macrophages to promote OXPHOS. In the body, macrophages exist in an environment of changing polarising cytokines, and so we sought to determine the effect of HBD3 when added to already polarised macrophages.

**Figure 5:**
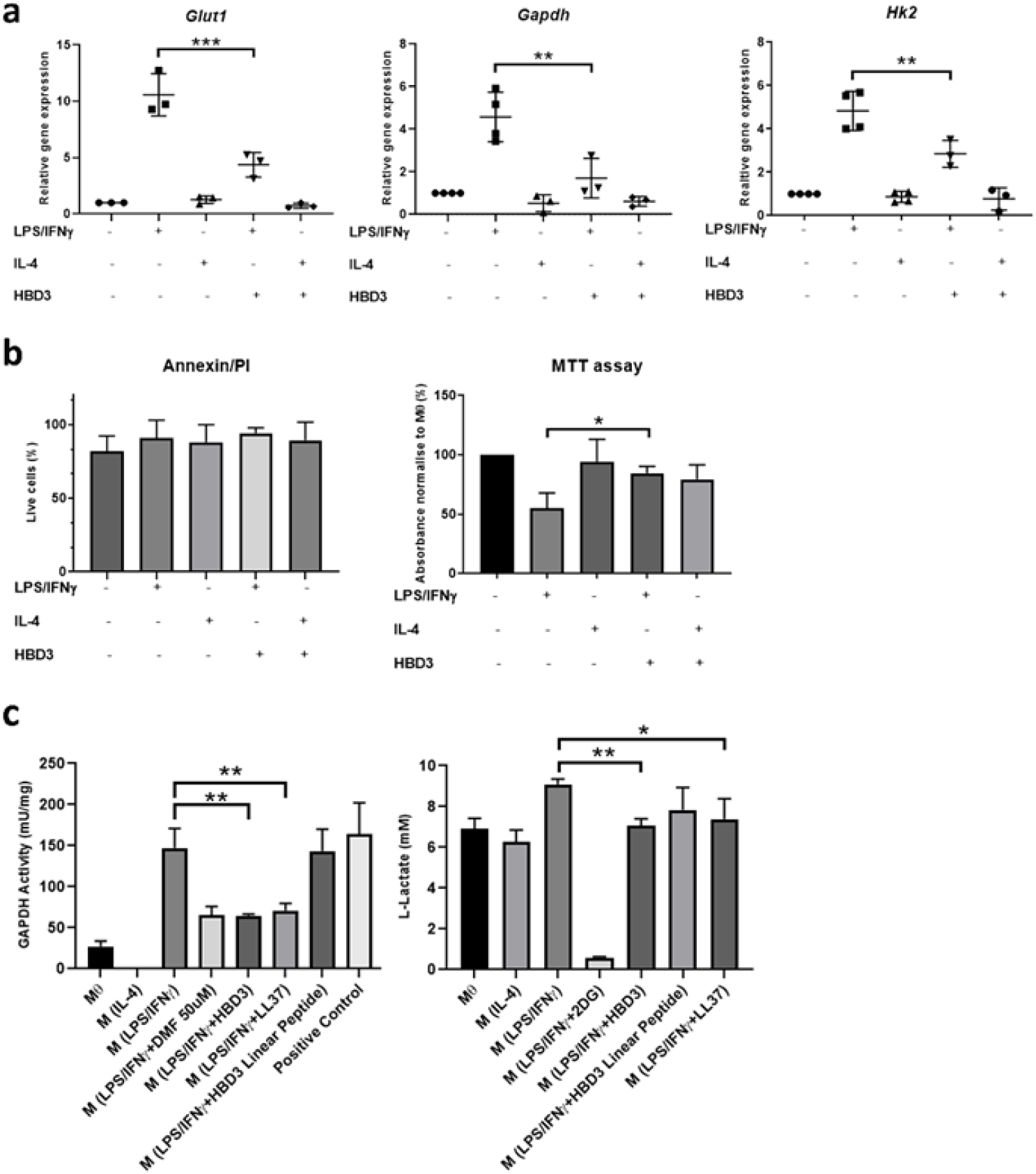
M(LPS+IFNγ)+HBD3 have lower glycolytic gene expression and GAPDH activity than M(LPS+IFNγ). **a:** *Glut1, Gapdh* and *Hk2* gene expression was measured by real time. Results are normalized to mRNA expression of naïve macrophages. *** p<0.001, ** p<0.01. **b:** Cell viability and metabolic activity measured respectively by Annexin/PI staining and MTT assay. The value for each sample was compared to naïve macrophages. * p<0.05. **c:** Glycolytic GAPDH activity and L-Lactate intracellular quantification assay respectively. ** p<0.01, * p<0.05. Comparison were done with one-way ANOVA test.

### M(IL-4) re-polarised with LPS+IFNγ in the presence of HBD3, promotes OXPHOS respiration

BMDM treated with either LPS/IFNγ or IL-4 for 24 hours, were re-polarised with the opposing inducer(s), with or without HBD3 for a further 18 hours (**Fig 6a**). It has been shown previously that M(LPS/IFNγ) protected from repolarisation by the alternative activation stimulus of IL-4, and this it has been shown that this loss of metabolic plasticity is due to irretrievable mitochondrial dysfunction by ROS generation and nitric oxide production (Van den Bossche *et al*., 2016). In contrast, M(IL-4) can be fully re-polarised with LPS/IFNγ to no longer use OXPHOS. Our data replicated these findings (**Fig. 6b** and **Supplementary Fig. 5a&b**) and we found that after 24 hours, the presence of HBD3 did not enable IL-4 to restore OXPHOS in M(LPS/IFNγ) +(HBD3+IL-4) (**Fig. 6b**). M(IL-4)+(LPS/IFNγ) showed complete loss of OXPHOS but when HBD3 was added just before the LPS/IFNγ re-polarisation stimulus, a significant level of oxidative phosphorylation was promoted as indicated by increased basal, ATP and maximal respiration (**Fig. 6b** and **Supplementary Fig. 6a**).

**Figure 6:**
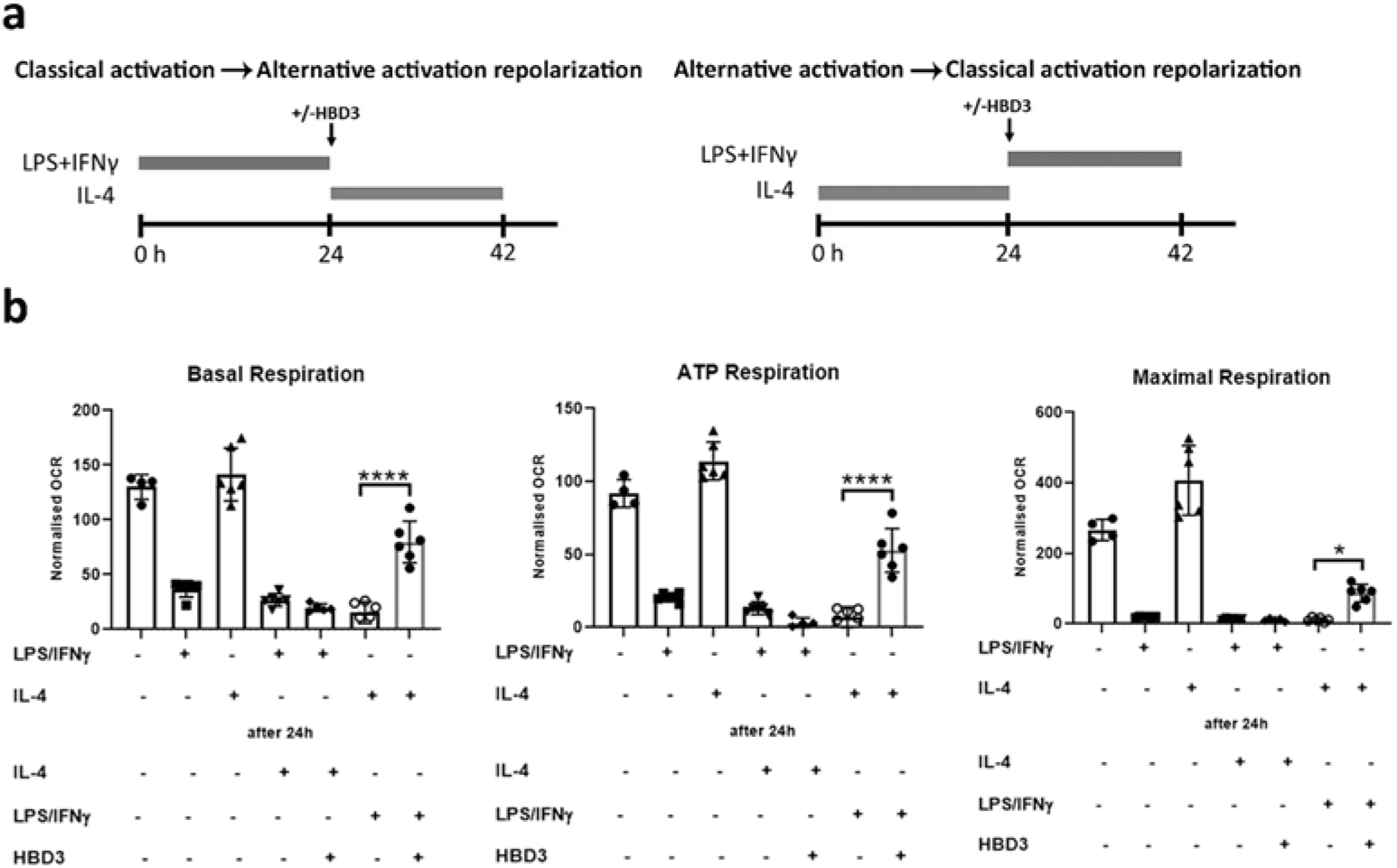
The presence of OXPHOS in M(IL-4) when re-polarised with LPS+IFNγ, requires the presence of HBD3. M(IL-4) or M(LPS/IFNγ) were treated with LPS/IFNγ or IL-4 respectively for 24 hours, with or without HBD3 for 18 hours. **a**: Schematic representation of re-polarisation condition. **Left Panel**: BMDMs were stimulates 24 hours with pro-inflammatory stimuli (LPS/IFNγ), followed +/-HBD3 for 30 minutes and then IL-4 for an additional 18 hours. **Right Panel**: BMDMs were stimulated for 24 hours with anti-inflammatory stimuli (IL-4), followed +/-HBD3 for 30 minutes then (LPS/IFNγ) for an additional 18 hours. **b:** Calculated basal respiration, ATP production and maximal respiration measured under mitochondrial stress condition. **** p<0.0001, * p<0.05. See **supplementary figure 6a** for representative experimental graph. Comparison were done with one-way ANOVA test.

Consistent with the presence of OXPHOS in M(IL-4)+(HBD3+LPS/IFNγ), the levels of both TNF-α and IL-6 were reduced and *Il4* gene expression was increased compared to M(IL-4)+(LPS/IFNγ) (**Fig. 7a**). Furthermore, CD86 and MHCII were expressed at significantly lower levels and CD206 and CD273 at higher levels in M(IL-4)+(HBD3+LPS/IFNγ) (**Fig. 7b & supplementary fig. 6b**). Thus, re-polarisation of M(IL-4) with LPS/IFNγ in the presence of HBD3, increased IL-4 production and limited pro-inflammatory activation and increased markers of alternative activation.

**Figure 7:**
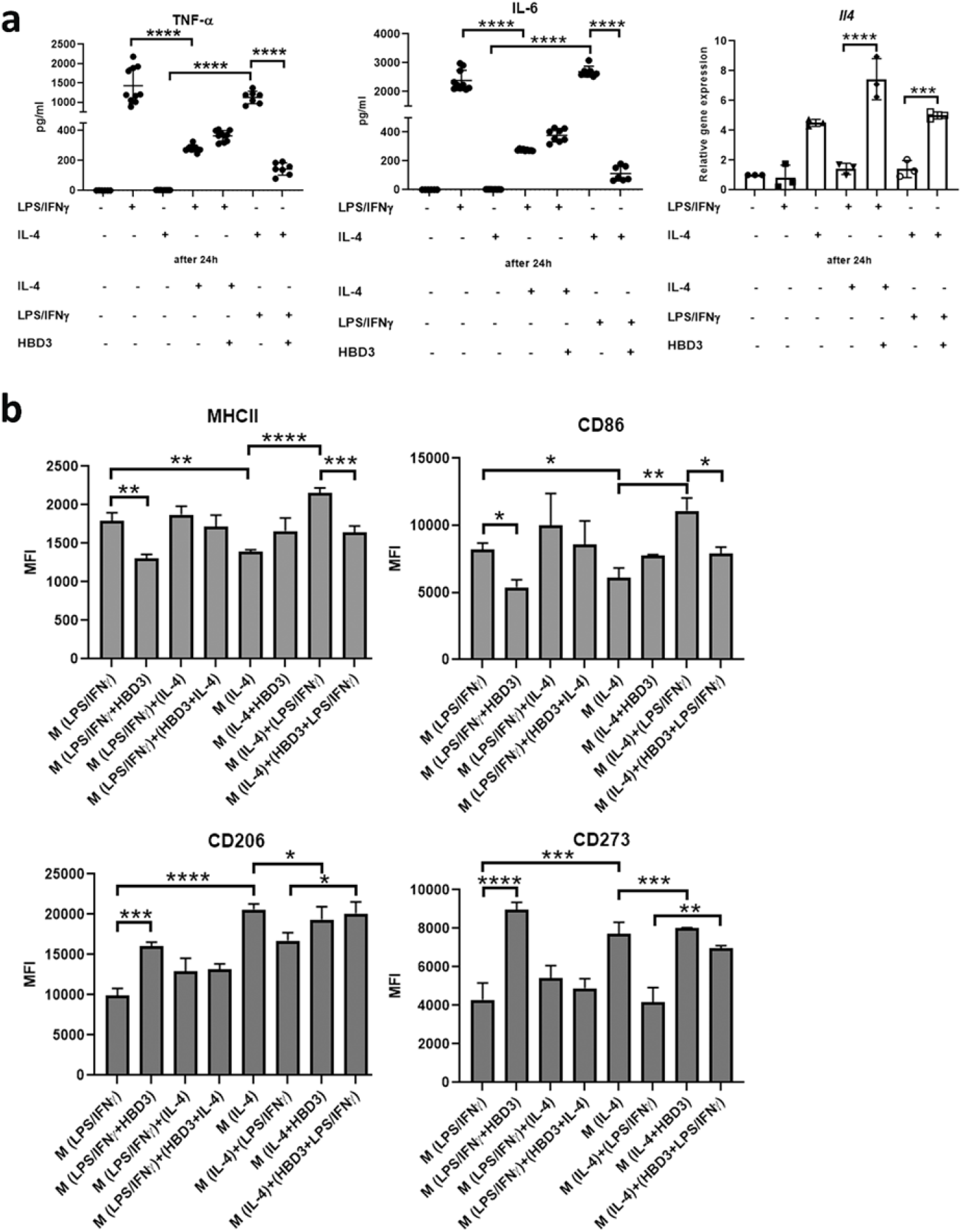
HBD3 limits M(IL-4) re-polarisation to an inflammatory phenotype. M(IL-4) or M(LPS+IFNγ) were treated with LPS+IFNγ or IL-4 respectively for 24 hours, with or without HBD3 for 18 hours. **a:** TNFα or IL-6 level and *Il4* gene expression was measured by ELISA and real time respectively. **** p<0.0001, *** p<0.001. **b**: Representative histograms of geometric mean fluorescence intensity (MFI) of MHCII, CD86, CD206 and CD273 surface markers analysed by flow cytometry. **** p<0.0001, *** p<0.001, ** p<0.01, * p<0.05. See **Supplementary figure S6** for corresponding dot plots

### HBD3 has a similar effect to IL-4 on the phenotype of MLPS/IFNγ

The significant increase in *Il4* expression observed when HBD3 was present in BMDM stimulation with LPS/IFNγ, promoted us to directly compare the effect of IL-4 and HBD3 on LPS/IFNγ macrophage polarisation. Macrophages were stimulated with LPS/IFNγ in the presence of different concentrations of IL4 (either 20 ng/ml (**Fig. 8**) or titrated doses from 0.5-20 ng/ml **(Supplementary Fig. 7**)), and compared to the effect of HBD3 with or without the cytokine. The reduction in expression levels of MHCII and CD86 observed when HBD3 was added after LPS/IFNγ, was similar to that seen when 20ng IL-4 was substituted for HBD3, and the effect was not additive for MHCII, but was for CD86 cell surface expression (**Fig. 8a**). The increase in MFI of CD206 in M(LPS/IFNγ+HBD3) compared to M(LPS/IFNγ) was also seen when IL-4 replaced HBD3, but 20ng M(IL-4) was more potent and 5μg/ml HBD3 had an equivalent effect to 0.5ng/ml IL-4 (**Supplementary Fig. 7**). This was mirrored by CD273, but here the combination of HBD3 and IL4 was additive. The level of TNF-α was found to be reduced in M(LPS+IFNγ) to the same degree by either HBD3 or IL-4, and the two together did not decrease the reduction level further. The effect on IL-6 was similar for HBD3 or 20ng/ml IL-4 when applied independently, but together the decreased response was additive **(Fig. 8b** and **Supplementary Fig. S7)**.

**Figure 8:**
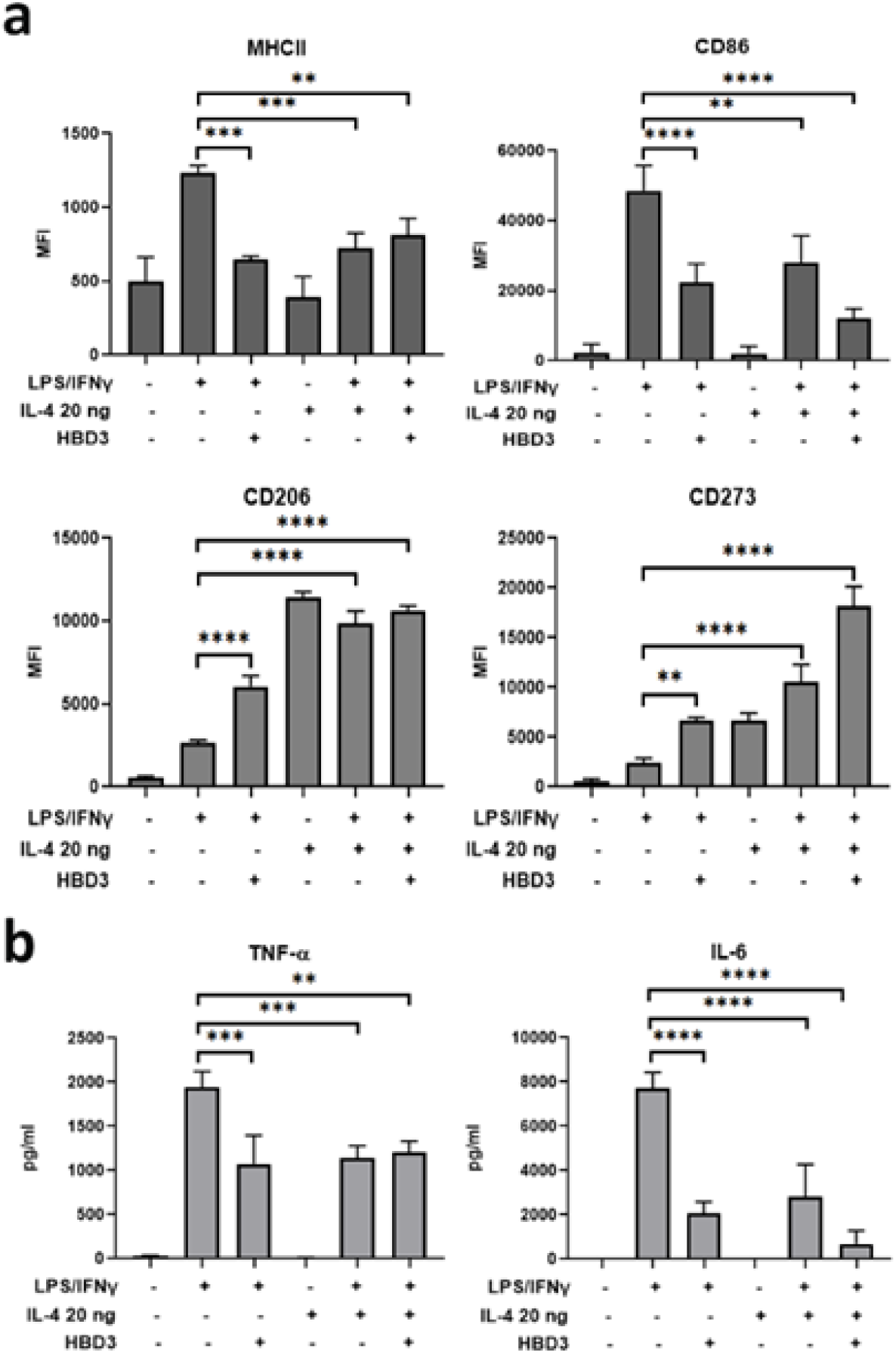
HBD3 has similar effect on M(LPS+IFNγ) as IL-4. BMDM were polarised with (LPS+IFNγ) or IL-4 20 ng, with or without HBD3 for 18 hours. **a**: Representative histograms of geometric mean fluorescence intensity (MFI) of MHCII, CD86, CD206 and CD273 surface markers analysed by flow cytometry. **** p<0.0001, *** p<0.001, ** p<0.01. **b:** TNF-α and IL-6 expression in supernatant measured by ELISA. **** p<0.0001, *** p<0.001, ** p<0.01. Comparison were done with one-way ANOVA test.

### HBD3 effect on M(LPS+IFNγ) polarisation is dependent on IL4-Receptor alpha

The HBD3-mediated suppression of classical activation markers and pro-inflammatory cytokines in M(LPS+IFNγ), and the production of *Il4*, prompted us to determine whether IL-4 was causative using BMDM from *Il4ra* knockout mice. The expression of *Il4* in M(LPS/IFNγ+HBD3) observed in WT mice, was lost in cells from *Il4ra*^-/-^ mice, demonstrating that the induction of *Il4* expression is dependent on IL4-Receptor alpha (**Fig. 9a**). In addition, the ability of HBD3 to reduce the expression of MHCII and CD86, and increase CD206 and CD273 in M(LPS+IFNγ), was lost in cells from *Il4ra* KO (**Fig. 9b**). *Arg1* and *Retnla* expression was also lost, as expected, in the *Il4ra* KO cells (**Supplementary Fig. 8**). TNF-α and IL-1β cytokine levels were increased by a comparable degree in both wild type and *Il4ra*^-/-^ M(LPS/IFNγ), but when HBD3 was included in the polarisation, the reduction in cytokine levels seen in wild type cells, was lost in *Il4ra*^-/-^ (**Fig. 9c**). In contrast, IL-6 levels in M(LPS/IFNγ+HBD3) were still reduced compared to the levels detected in M(LPS/IFNγ), independent of *Il4ra* expression (**Supplementary Fig. 8**). These data show that the suppressive effect of HBD3 on classical activation is primarily, but not exclusively, dependent on IL-4Rα.

**Figure 9:**
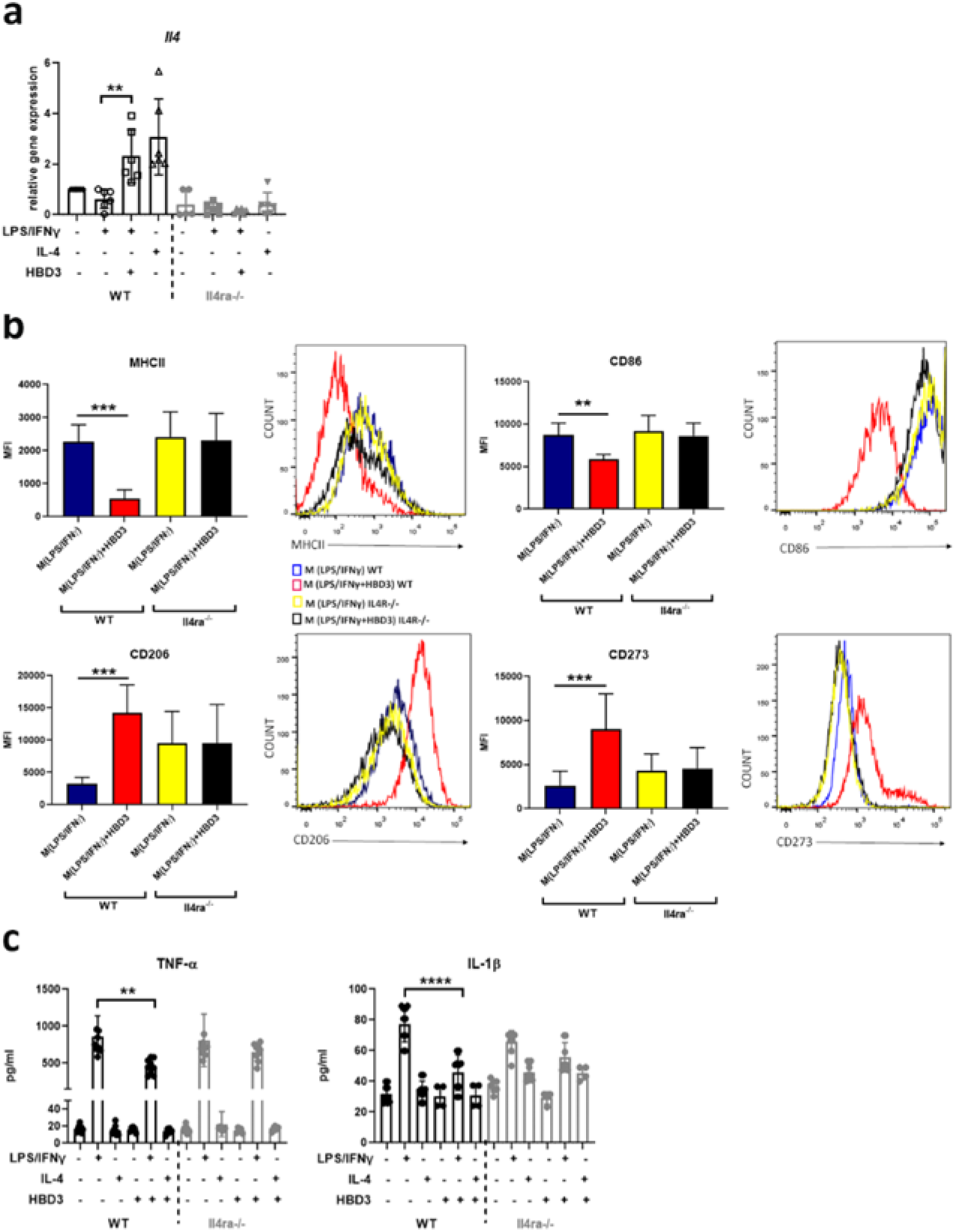
HBD3 effect on M(LPS+IFNγ) is dependent on IL-4Rα. BMDM were polarised with (LPS+IFNγ) or IL-4 20 ng, with or without HBD3 for 18 hours. **a:** *Il4* gene expression were measured by real time. Results are normalized to mRNA expression of naïve macrophages. ** p<0.01. **b:** Representative histograms of geometric mean fluorescence intensity (MFI) of MHCII, CD86, CD206 and CD273 surface markers analysed by flow cytometry. *** p<0.001, ** p<0.01. **c:** TNF-α and IL-1β expression in supernatant was measured by ELISA. **** p<0.0001, ** p<0.01. Comparison were done with one-way ANOVA test.

## Discussion

Defensins are potent antimicrobials predominantly secreted from epithelia at outward facing surfaces. However, their diverse immunmodulatory actions to increase or decrease inflammatory responses implies that they are involved in other processes *in vivo*. We show here a novel function for the antimicrobial peptide, β-defensin HBD3 (DEFB14 in mouse), in suppression of the classical activation response of macrophages, and increase in alternative activation. *Defb14* ^*tm1a*^ mice displayed an increased inflammatory serum cytokine in the acute response to LPS. *In vitro*, polarisation with the proinflammatory cocktail (LPS/IFNγ) and subsequent addition of HBD3, resulted in human and mouse macrophages, that show decreased inflammatory cytokine secretion and expression of IL-4, and an increase in M(IL-4) alternative activation markers.

Consistent with the decrease in classical activation and increase in alternative activation there was a change in metabolic flux, with an increase in OXPHOS and a reduction in genes key for aerobic glycolysis.Cellular metabolic re-programming is recognised as an important component in activation-induced inflammation or homeostasis (Murphy, 2019; Viola *et al*., 2019).

The functional switch of macrophages to a pro-inflammatory, protective but potentially host-damaging phenotype, is self-limited through a variety of mechanisms that include lactylation (Zhang *et al*, 2019), production of anti-inflammatory molecules such as IL-10 (Ip *et al*., 2017), and itaconate -which relies on a break in TCA cycle (Lampropoulou *et al*, 2016). Lactate increases during glycolysis and has been shown to mediate lactylation of histone lysines (Kla) in promoter regions of core genes, 16 to 24 hours after M1 activation (Zhang *et al*., 2019). Lactate supplementation can increase histone Kla levels, and induces *Arg1* expression through Kla of the *Arg1* promoter. It is unlikely that lactylation is relevant here as consistent with a reduced glycolytic flux in M(LPS/IFNγ) in the presence of HBD3, lactate levels decreased indicating no promotion of lactylation by the presence of HBD3 at 18 hours, despite increased markers of M(IL-4) alternative activation. In addition, we show here that HBD3 enhances the IL-4 mediated polarisation of macrophages, a process which does not involve lactylation(Zhang *et al*., 2019).

A shift in metabolic flux that promotes OXPHOS has been shown to be mediated by IL-10 in macrophages exposed to LPS, with inhibition of glucose uptake and glycolysis (Ip *et al*., 2017). We did see that M(LPS/IFNγ+HBD3) had a high IL-10 level at 18 hours and increased OXPHOS, but the IL-10 level was less than that evident in M(LPS/IFNγ). Thus, IL-10 is likely not central to the ability of HBD3 to increase OXPHOS.

Activated macrophages and lymphoid cells, can be succinated by DMF to inactivate the catalytic cysteine of GAPDH, to reduce glycolytic flux both *in vitro* and *in vivo* (Kornberg *et al*, 2018). A similar reduction in GAPDH activity in M(LPS/IFNγ) was seen here when we substituted 5μM HBD3 for 50μM DMF. The antimicrobial peptide cathelicidin, can also reduce LPS-induced cytokine response in macrophages (Hancock *et al*, 2016; Sun *et al*, 2015). Although cathelicidin has a simple linear alpha helical structure, unlike the conserved cysteine knot structure of defensins, in common with HBD3 it is amphipathic and cationic, and can rapidly enter macrophages. LL-37 has been shown to associate with GAPDH in macrophages (Mookherjee *et al*, 2009). It is possible that HBD3 binds GAPDH to inhibit its activity and limit expression of key enzymes of glycolysis, and reduce production of lactate. A consequence of this would be that glycolysis would stall at glyceraldehyde 3 phosphate, but the pentose pathway would remain intact, allowing N-glycosylation, essential for CD206 cell surface expression, which was significantly increased by HBD3. LPS induction of IL-1β (but not TNF-α) has been shown to be dependent on succinate accumulation during glycolysis (Tannahill *et al*, 2013). HBD3 reduces both these cycokines and so does DMF(Ali *et al*, 2020).

In addition to reduction in the macrophage polarisation reponse to LPS/IFNγ, we also show here that, mouse BMDM, polarised with sub-saturation levels of IL-4, show an augmented alternative activation when in the presence of HBD3. When HBD3 is present duinrg the LPS/IFNγ polarisation, IL-4 gene expression is induced in BMDM, RAW 264.7 cells and cytokine is found in PBMDM. IL-4 and HBD3 had a similar effect on the phenotype of cells exposed to LPS/IFNγ and the effect on IL-6 secretion was additive even at high concentration of IL-4. Macrophages activated by IL-4, can induce transcriptional suppression of a subset of genes that results in reduced responsiveness to LPS by a subset of pro-inflammatory genes including IL-1β (Czimmerer *et al*., 2018). It is likely therefore that HBD3 driving IL-4 production is key in the effects of the peptide on polarisation. Several publications have previously reported IL-4 production by macrophages under certain conditions. The autocrine production of IL-4 by macrophages in response to TLR activation was found at 24-48 hours (Mukherjee *et al*, 2009; Shirey *et al*, 2010). In addition, IFNγ primed BMDM exposed to LPS in combination with immune complexes that ligate FcγRs, have also been shown to potently induce production of IL-4 cytokine (La Flamme *et al*, 2012).

The ability of HBD3 to decrease the expression of classical activation markers and cytokines TNF-α and Il-1β and increase alternative activation markers in M(LPS/IFNγ) was dependent on IL-4Rα. This again indicates the significance of the observed increase in IL-4 in the presence of classical stimulation and HBD3. HBD3 mediated reduction of IL-6 secretion however, was not dependent on IL-4Rα. Thus HBD3 must act through IL-4Rα dependent and independent mechanisms.

It is important to consider whether the immunomodulatory effect of HBD3 is likely to be relevant *in vivo*. The inability of HBD3 to promote OXPHOS in cells, already polarised for 24 hours with LPS/IFNγ, is similar to the findings using IL-4 *in vitro* and also *in vivo* (Van den Bossche *et al*., 2016). Ruckerl et al. demonstrate macrophage plasticity is present *in vivo*, and the type II activation phenotype established due to an initial nematode infection, can be altered upon subsequent infection with *Salmonella* (Rückerl *et al*, 2017). In keeping with *in vitro* results, there was no evidence that the *Salmonella activation* can be altered by subsequent *nematode* challenge. However DMF, as discussed above, is an effective treatment for multiple sclerosis and psoriasis as an immunomodulatory compound, and DMF alters the metabolic flux in macrophages and lymphocytes to reduce glycolysis and influence survival of Th1 and Th17 cells(Kornberg *et al*., 2018).

The fact that HBD3 reduced production of damaging inflammatory cytokines (including IL-12 which influences both innate and adaptive responses) is encouraging that a change in myeloid metabolism from glycolytic to OXPHOS may be relevant *in vivo* if given rapidly or chronically. DEFB14 and HBD3 peptides have been shown to induce polarisation of CD4+ T helper cells to *Foxp3* expressing CD4+ T cells, and can rescue mice from experimental acute encephalitis, a model of multiple sclerosis (Bruhs *et al*, 2016; Navid *et al*, 2012).

Several other lines of evidence support HBD3/DEFB14 being important in resolution and restoration of homeostasis. In addition to the *Defb14* ^*tm1a*^ response to LPS, we describe here, *Defb14* gene targeted mice show a global delay in wound healing *in vivo* (Williams *et al*, 2018). In addition, DEFB14 production from mouse pancreatic endocrine cells, stimulates IL-4 secreting B cells through TLR2 to increase alternate activation of macrophages, allowing tolerance and prevention of autoimimmune diabetes (Miani *et al*, 2018). Finally, Tewary et al. report that splenocytes from mice, immunised with OVA+/-HBD3, and isolated a week after a final boost, show an increase in the type 2 cytokine IL-5 when HBD3 was present (Tewary *et al*., 2013). HBD3 is, however, a double edged sword and can act both as an alarmin and a suppressor of inflammation. Indeed Tewary et al. also report OVA+CpG+HBD3 immmunisation induces cells expressing IFNγ. The difference in effects is likely due to the prevailing environment and the concentration of the peptide.HBD3 is one of the seven copy number variable β-defensins, and increased copy number and expression level associates with the inflammatory Th1/Th17 autoimmune disease psoriasis (Abu *et al*, 2009). The levels of HBD3 observed in the skin are The anti-inflammatory effect we describe here is somewhat of a paradox to this association, but a “goldilocks” effect may exist where too little or too much is not the right amount. Our titration of HBD3 showed lower levels of peptide suppress TNF-α production but this is diminished at high peptide levels that are known to be toxic (Leelakanok *et al*., 2015). Expression of β-defensins is normally low, around 0.2ng/ml in normal serum but inflammation can drammatically increase expression by up to 1000 fold in the serum of patients with the inflammatory condition psoriasis and even higher in the skin(Jansen *et al*, 2009).

In conclusion, we show here that HBD3 strongly influences macrophage polarisation through an IL-4Rα dependent mechansim. The data indicate that HBD3 drives IL-4 production under inflammatory conditions resulting in a limit on classical activation and pro-inflammatory responses *in vitro* and *in vivo* and coincident with augmentation of macrophage alternative activation. HBD3 is rapidly induced by exposure to pathogen moelcular patterns or inflammatory cytokines and secreted from epithelial cells where it can act as a potent antimicrobial, but once the danger is reduced, its immunomodulatory properties may be a key part of the resolution process. Further work is required to investigate the *in vivo* ability of HBD3 or derivatives to control both innate and adaptive immune phenotypes and ascertain their potential for clinical and therapeutic benefit.

## Methods

### In vivo experiments

ES cell *Defb14*^*tm1a(HGU1)*^ gene targeted mice were generated as detailed previously (Reijns *et al*, 2012) using a KOMP vector and further information is provided in supplementary figure 1. Animal studies were covered by a Project License, granted by the UK Home Office under the Animal Scientific Procedures Act 1986, and locally approved by the University of Edinburgh Ethical Review Committee. Chimaeric mice were backcrossed to 8 generations on C57Bl/6J. Mice were injected intraperitoneally (IP) into male mice (8-12 weeks old) as previously described with 15mg/Kg LPS +/-10μg HBD3 peptide(Semple *et al*., 2010).

### Isolation, differentiation and polarisation of Macrophages

Mouse bone marrow cells were collected from the femurs and tibia of 6-10 weeks old C57BL/6 mice. After washing in DPBS (GIBCO™, #14040-091), cells were cultured at 2×10^6^ cells/plate in DMEM/F12 GlutaMAX™(GIBCO™, #31331-028) medium supplemented with 10% FBS (GIBCO™, #10500-064), 1% L-Glutamine (200 nM) (GIBCO™, #25030-024), 1% Penicillin-Streptomycin (10,000 U/mL) (GIBCO™, #15140-122) and 20% of L929-conditioned media, for 7 days. L929 conditioned media was made as previously reported (Weischenfeldt and Porse, 2008).5×10^5^ L929 cells were seeded in a T75 flask in 25ml of DMEM/F12 GlutaMAX™medium supplemented with 10% FBS, 1% L-Glutamine, 1% Pen-Strep. Cells were cultured for 7/8 days and the culture supernatant was collected, centrifuged for 5 minutes at 1200 rpm and then stored at 80°C. The cells were maintained in a humidified incubator with 95% air and 5% CO_2_ atmosphere at 37°C. The media was replaced every 2-3 days during the culture. After differentiation all the medium was removed, and naïve macrophages were stimulated to a classical, PRO-inflammatory phenotype (M(LPS+IFNγ) by overnight incubation with LPS (50 ng/ml).

Peripheral blood mononuclear cells were isolated by Percoll gradient from whole blood donated by healthy volunteers with written informed consent (AMREC 20-HV-069). Cells were plated at 2 x10e6 in 24-well tissue culture plates and cultured in RPMI + 10% FCS (low endotoxin), and matured to monocyte derived macrophages (MDMs) over 14 days, with media changes every 3 – 4 days. MDMs were then treated for polarisation as described in the text. Following treatment, cell supernatant was harvested for ELISA. (Lipopolysaccharides from Escherichia coli O111:B4, ultrapure, Source Bio Science, #AV-7016-1) and recombinant murine IFNγ (20 ng/m(PEPROTECH, #315-05)) or towards an alternative activation phenotype (M(Il-4)) by overnight incubation with recombinant murine IL-4 (20 ng/ml or other dilution as described in results (PEPROTECH, #214-14).

Treatments with HBD3 peptide(s) was after 15-30 minutes 5 μg/ml of human β-Defensin-3 (hBD3) (Peptide Institute Inc., PeptaNova GmbH #4382-s), or 5 μg/ml of Linear Defensin (Almac Sciences Scotland Ltd, UK) or 5 μg/ml of LL37 (Almac Sciences Scotland Ltd, UK). After 18 hours, medium and cells were harvested. For GAPDDH activity assay, cells were treated with Dimethyl fumarate (DMF) 50 μM (Sigma, #242926) and for Glycolysis Cell-Based assay, cells were treated with 2-deoxyglucose (2-DG) 100 nM (Sigma, #D8375).

### Flow cytometry analysis

Cells were cultured and treated as previously described. After washing, they were incubated with blocking solution (0.5% BSA, 1% FBS) for 10 minutes at room temperature. Then cells were incubated with: F4/80 (Biolegend, #123113), CD11b (BioLegend, #101261), MHCII (BioLegend, #107643), CD86 (BioLegend, #105041), CD206 (BioLegend, #141707), CD273 (BioLegend, #107205) antibodies for 30 minutes on ice. Samples were analysed on Flow Analyser QFCF 5L LSR FORTESSA (BD Bioscences), followed by data analysis using FlowJo (version X) flow cytometry analysis software (FlowJo, LLC). Isotype control antibodies used at the same concentration did not give any detectable signal. Data are representative of three independent experiments.

### RNA extraction and qRT-PCR

Total RNA from BMDMs was extracted with TRIzol™ Reagent (ThermoFisher Scientific, #15596026), followed by phenol:chloroform extraction with an overnight isopropanol precipitation. RNA samples were treated with 1 µl DNase I (2 U) (Ambion™, #AM2222) per 10 μg of total RNA in a 50 μl reaction for 30 minutes at 37°C, and then precipitated in 70% isopropanol with sodium acetate 150 mM. RNA concentration was determined by NanoDrop 1000 Spectrophotometer (ThermoScientific).1 µg of total RNA was reverse transcribed into cDNA with M-MLV Reverse Transcriptase 200 units (Promega, #M1701), Random Primer 0.5 μg (Promega, #C1181), dNTP Mix 200 μM (Promega, #U1511), RNasin® Ribonuclease Inhibitors 25 units (Promega, #N2511). Samples were incubated for 1 hour at 37°C. The cDNA generated was used for semi quantitative PCR on a StepOne plus RT PCR 96 well cycler (Applied Biosystem) according to the manufacturer’s instructions using Fast SYBR™ Green Master Mix (ThermoFisher, #4385616). Each PCR series included a no-template control that contained water instead of cDNA and a reverse transcriptase-negative control. Data was analysed on StepOne™Software v2.3 (Applied Biosystem) and target gene expression was normalized to the expression of the housekeeping gene ACTIN BETA. Relative gene expression was calculated using the standard 2-ΔΔCT method. Primers were designed using PrimerQuest Tool (IDT Integrated DNA Technologies). Samples were analysed in technical triplicates and biological triplicates.

### ELISA

Levels of TNF-α, IL-6, IL-1β, IL-10, IL-4 in cell culture supernatants or peritoneal washes were determined using DuoSet ELISA Development kits (R&D Systems) according to the manufacturer’s instructions. Samples were analysed in technical duplicates and biological triplicates, each standard in technical duplicate.

### Measurement of Oxygen Consumption Rate (OCR)

The OCR was measured using a Seahorse XFe24 Analyzer (Seahorse Biosciences, Billerica, MA, USA). BMBMs were seeded in XF24 cell culture microplates (24 wells) at a density of 2 × 10^5^ cells/well and incubate overnight. After treatment as described before, assay medium consistent of Seahorse XF Assay medium (Agilent, Seahorse Bioscences, #102365-100) supplemented with Glucose 10 mM (Sigma, #G8270), Sodium pyruvate 2 mM (Sigma, #P5280) and adjusted to pH 7.4, was added to the cells. The inhibitors and uncouplers used in this study were as follows: Oligomycin A 2μm (Sigma, #75351), FCCP 75 μM (Cambridge Bioscience, #15218), Rotenone 1 μM (Sigma, #R8875) and Antimycin A 2.5 μM (Sigma, #A8674). OCR was normalised to cell number by SRB staining. Each sample was assayed in 6 technical replicates, and 2 biological replicates.

### Sulforhodamine B (SRB) staining

Cells seeded in a 24 well plate in presence of medium, were fixed with 50% TCA solution (Sigma, Trichloroacetic acid #T9159) at the final concentration of 10%, and incubate at 4°C for 1 hour. After wash in tap water and air-dried, TCA-fixed cells were stained by adding 50 μl of SRB solution 0.4% in 1% glacial acetic acid (Sigma, SRB #S1402) for 30 minutes at room temperature. The excess dye was removed with 10 washes with 2% glacial acetic acid (Fisher Chemical, #A38S-500). The plate was air-dried and the cell-bound dye re-dissolved by 100 μl of Tris solution 10 mM pH 10.5 (Sigma, Trizma #T6066). The OD was measured at 540-490 nm with a microplate reader (SYNERGY™ HT, BIOTEK® Instrument Inc., Vermont USA).

### GAPDH assay

Glyceraldehyde 3 Phosphate Dehydrogenase Activity was measured by using Glyceraldehyde 3 Phosphate Dehydrogenase Activity Assay kit (Colorimetric) (Abcam®, #ab204732) according to the manufacturer’s instructions. Briefly, cells were cultured in 6 wells plate and treated at different condition. DMF was added as a negative control. After lysis, the supernatant was collect and added to 50 μl of Reaction mix. The OD of standards, samples and positive control was measured at 450 nm, in a microplate reader in a kinetic mode, every 2 minutes, for 10 minutes. Data were analysed as the kit’s manual suggested. Each sample was assayed in technical triplicate and biological triplicates, and each standard in technical duplicate.

### Lactate assay

Glycolysis Cell-Base Assay kit (Cayman Chemical, #600450) was a colorimetric method to detect L-lactate amount in the medium. According to the manufacturer’s instructions, cells seeded in a 96 wells plate, were treated and grown one overnight in absence of serum. As negative control cells were treated with 2-2DG. 10 μl of medium for each sample and 10 μl of each standard were added to reaction solution and incubated for 30 minutes in an orbital shaker at room temperature. The absorbance was measured at 490 nm in a microplate reader. Data were analysed as the kit’s manual suggested. Each sample was assayed in technical triplicate and biological triplicates, and each standard in technical duplicate.

### MTT assay

Succinate dehydrogenase activity was measured by MTT assay (Sigma, #M2128). Briefly, BMDMs were seeded in a 96 well plate and incubated with 100 μl of MTT 5mg/ml at 37°C. After 3 hours the media was removed and acidic isopropanol (0.1 N HCl) was added for 30 minutes. The succinate dehydrogenase activity was assessed by yellow MTT reduction into purple formazan and the absorbance was measured at 580 nm in a microplate reader. Each sample was compared to the level of naïve macrophages, and was analysed in technical triplicates and biological triplicates.

### Cell viability

Cell viability was tested by using the APC Annexin V Apoptosis Detection Kit (BioLegend, #640932). Cells were washed twice with cold BioLegend Cell Staining Buffer, and then re-suspended in Annexin V Binding Buffer at a concentration of 0.25-1.0 × 10^7^ cells/ml. Cells were stained with 5 µl of APC Annexin V and 10 µl of Propidium Iodide Solution for 15 min at room temperature in the dark. The samples was analysed by flow cytometry.

### Re-polarisation

Cells was treated with LPS/IFNγ or IL-4 and after 20/24 hours washed in PBS, and then treated with medium. After 30 minutes, polarisation factors were added as indicated. For HBD3 treatment, HBD3 was added 30 minutes before the re-polarisation factors.

### Statistical analysis

Statistical analyses were performed using GraphPad Prism 8.1.2 (GraphPad Software Inc., San Diego). Comparison were done with one-way ANOVA test, * p<0.05, ** p<0.01, *** p<0.001, **** p<0.0001. Statistical values can be found in the figure legends.

## Acknowledgements

Flow cytometry data was generated with support from the QMRI Flow Cytometry and cell sorting facility, University of Edinburgh. We thank Drs. Dietmar Zaiss and Pieter Louwe for valuable discussions. We would also like to thank staff at the BVS for expert technical assistance. This work was supported by MRC Human Genetics Unit core award to JRD 2010-15 and Medical Research Council UK grant (MR/P02338X/1) awarded to JRD and NMN. BJM supported by MRC SHIELD consortium MR/N02995X/1 grant award to DJD.

M.E.C. designed and performed most experiments, analysed and interpreted the data,. D.J.P.A performed experiments and contributed to discussion, R.N.C. assisted and oversaw Seahorse experiments and analysed data F.S. performed experiments and contributed to discussion, F.K. performed ES cell targeting, S.W. performed experiments, D.T. assisted in analysis of Flow Cytometry data, H.J.W.M. carried out the human cell experiments, B.J.M. supervised human cell experiments and analysis, D.H.D. supervised the human experiments and contributed to design, D.J.D. and J.E.A. provided critical feedback, S.J.J. assisted with data interpretation and critiqued the manuscript for intellectual content. N.M. and J.R.D. conceived, designed, and supervised the project, JRD interpreted the data and wrote the manuscript.

