## supplemental files for "Classical macrophage polarisation is limited by human β-defensin-3 via an autocrine IL-4 dependent process" (from another version of this preprint: version 1, 2021-05-06)

**Supplementary Figure 1**

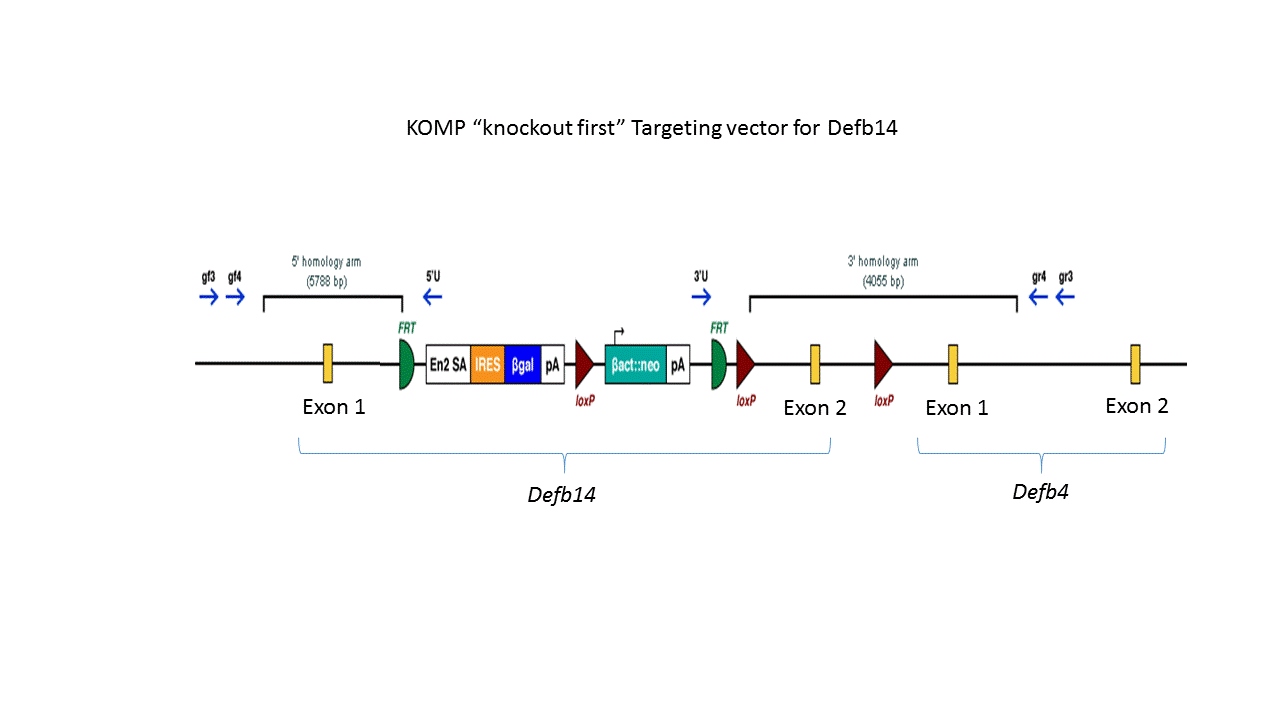

**Vectors for creating Defb14 knockout first mice.**

Image to show the tm1a targeting vector illustrating how the integration creates a disrupted gene between exon 1 and 2. The vector [CSD49720](http://www.mousephenotype.org/data/search?term=Defb14&type=gene) was generated by the trans-NIH Knock-Out Mouse Project (KOMP) and obtained from the KOMP Repository ([www.komp.org)*](http://www.komp.org)*). Mutation was created in house in E14IV Ola 129/Ola ES cells (obtained as kind gift from Prof. Austin Smith) by electroporation as previously described (Reijns et al. 2012). Mice were backcrossed to 8 generations onto C57Bl/6N and bred as heterozygotes. Homozygotes appeared healthy without obvious phenotype and both males and females were fertile**.**

**NIH grants to Velocigene at Regeneron Inc (U01HG004085) and the CSD Consortium (U01HG004080) funded the generation of gene-targeted ES cells for 8500 genes in the KOMP Program and archived and distributed by the KOMP Repository at UC Davis and CHORI (U42RR024244). For more information or to obtain KOMP products go to www.komp.orgobtained from Knockout mouse project KOMP )*

**Supplementary Figure 2**

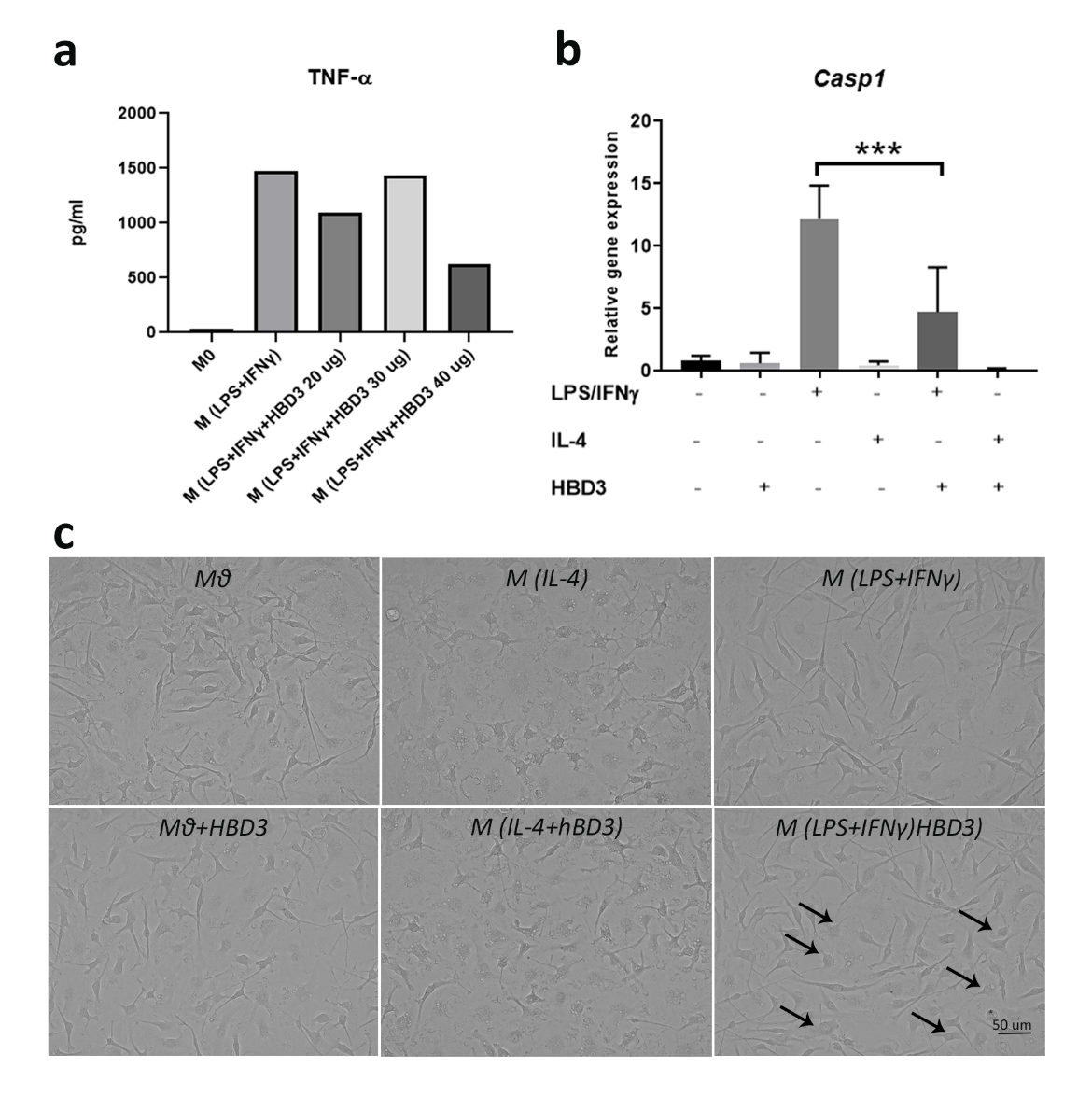

**HBD3 change pro-inflammatory macrophages morphology**

**a:** TNF-α expression in supernatant of BMDMs Mθ and M(LPS+IFNγ) treated with different HBD3 concentration (20, 30, 40 ug) for 18 hours, measured by ELISA. **b:** Casp1 gene expression measured by real time in BMDM polarised as shown with or without (5 ug/ml) HBD3 for 18 hours, *** p<0.001. Comparison were done with one-way ANOVA test. **c:** Morphology of BMDM under various polarisation conditions. Arrows indicate cells with morphology different to that seen in M(LPS+IFNγ).

**Supplementary Figure 3**

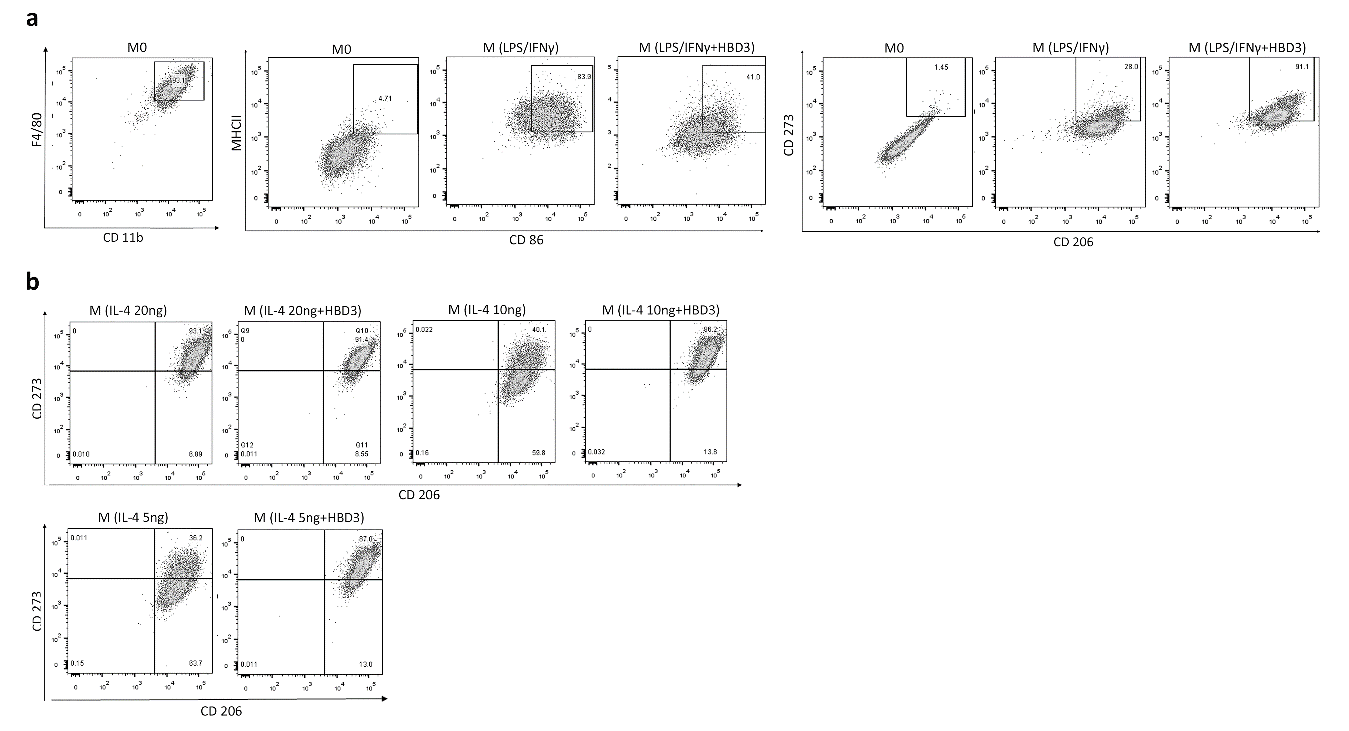

**HBD3 decreased and increased respectively pro-inflammatory and alternative activation markers in M (LPS+IFNγ), besides augments IL-4 activation.**

**a:** Dot plots of the different patterns of expression of F480/CD11b, MHCII/CD86 and CD273/CD206 surface markers for data shown in Figure 2a. The dot plots in row A show: 83.9% double positive MHCII/CD86 and 28% double positive CD273/CD206 population observed in M (LPS/IFNγ), compared to 41% double positive MHCII/CD86 and 91.1% double positive CD273/CD206 population observed in M (LPS/IFNγ+HBD3).

**b:** Dot plots of the different patterns of expression of CD273/CD206 surface markers for data shown in Figure 3b. Double positive CD273/CD206 populations observed respectively in M (IL4 20ng); M (IL4 20ng+HBD3); M (IL4 10ng); M (IL4 10ng+HBD3); M (IL4 5ng); M (IL4 5ng+HBD3) were 93.1%; 91.4%; 40.1%; 86.2%; 36.2; and 87%. Debris and aggregates were eliminated from the analysis by forward- and side-scatter analysis, isotype controls and compensation protocols were used.

**Supplementary Figure 4**

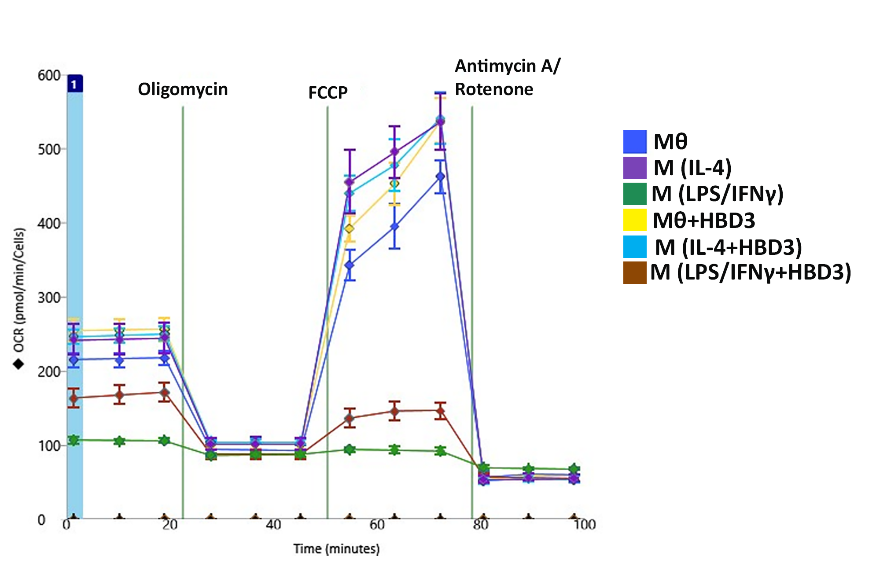

**Mitochondrial respiration in M(LPS/IFNγ+HBD3)**

Representative experiment of cellular respiration measured under mitochondrial stress condition in M0, M (LPS/IFNγ), M (IL-4), with and without HBD3. Oligomycin inhibits ATP synthase (complex V), Carbonyl cyanide 4-(trifluoromethoxy)phenylhydrazone (FCCP) uncouples mitochondrial respiration and allows oxygen consumption by complex IV to reach its maximum capacity. Antimycin A/Rotenone block mitochondrial complex III and I and indicates non-mitochondrial oxygen consumption.

**Supplementary Figure 5**

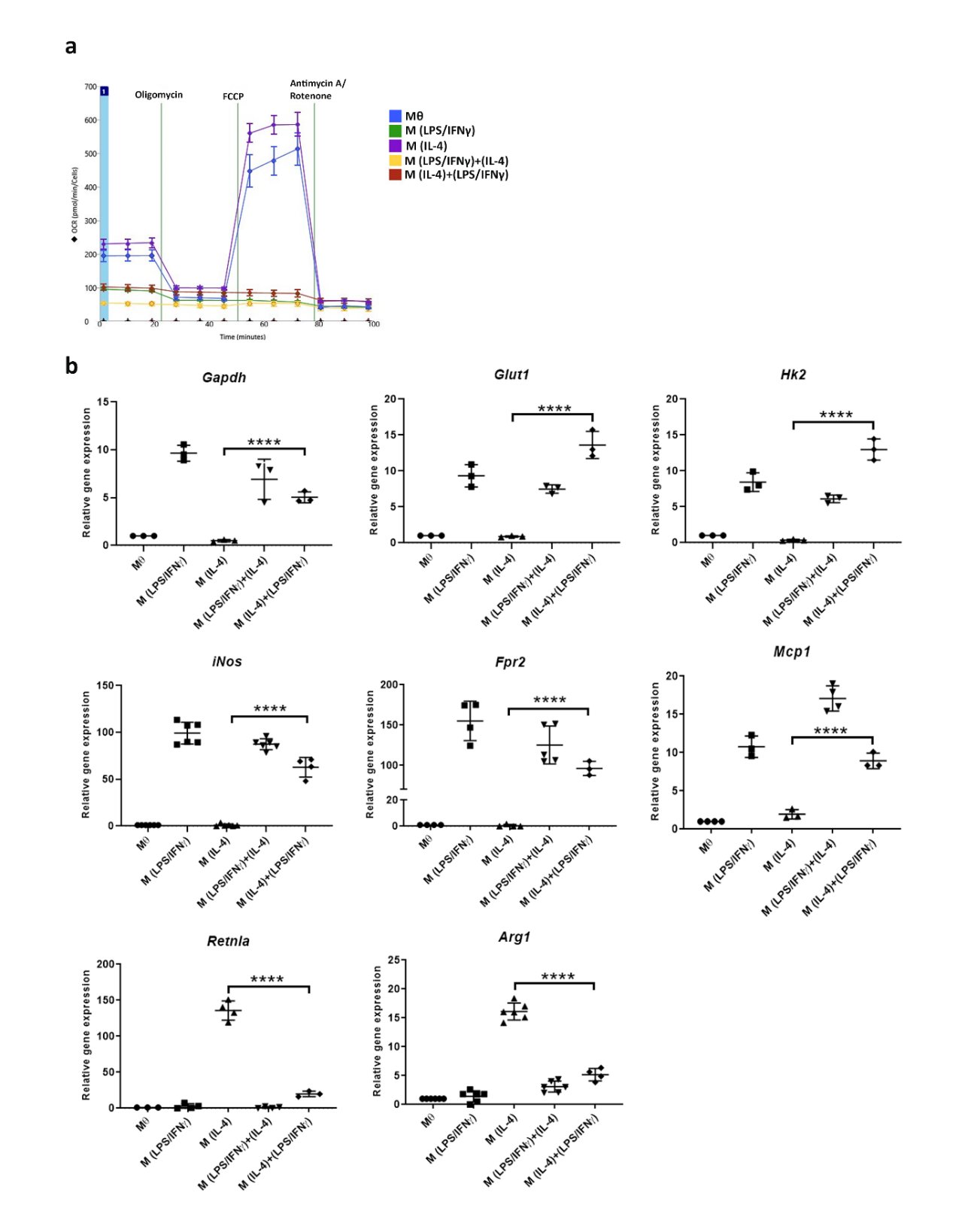

**M(LPS+IFNγ) re-polarised with IL-4 do not promote OXPHOS, but M(IL-4) re-polarised with LPS/IFNγ undergo metabolic re-programming.**

**a:** Representative experiment of cellular respiration measured under mitochondrial stress condition in cells repolarised as indicate. M0, M (LPS/IFNγ), M (IL4), M (LPS/IFNγ)+(IL4), M (IL4)+(LPS/IFNγ).

**b:** *Gapdh*, *Glut1*, *Hk2*, *iNOS*, *Fpr2*, *Mcp1*, *Retnla*, *Arg1* gene expression was measured by real time in the same samples. Results are normalized to mRNA expression of naïve macrophages. **** p<0.0001. Comparison were done with one-way ANOVA test.

**Supplementary Figure 6**

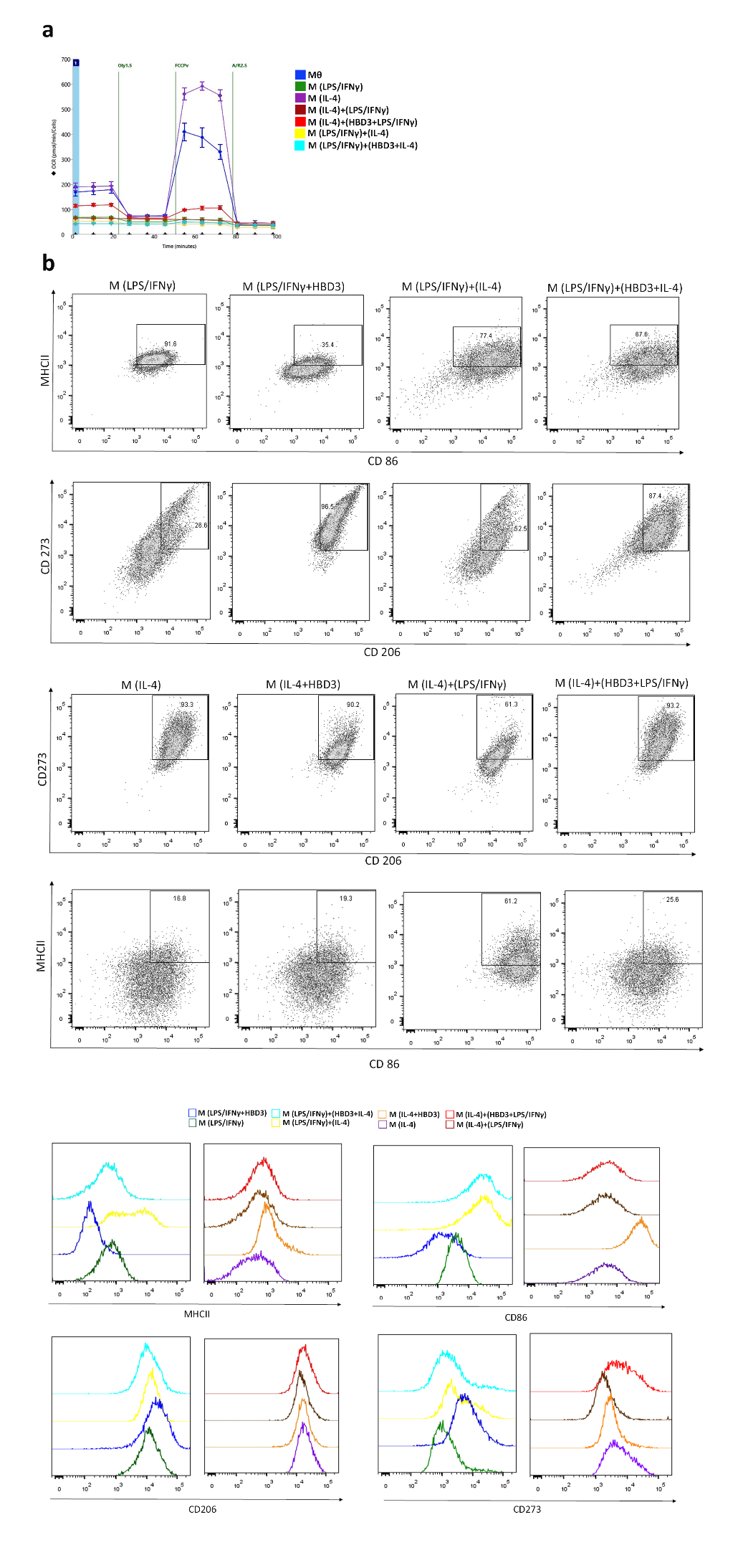

**HBD3 limits M(IL-4) re-polarisation to an inflammatory phenotype.**

**a:** Representative experiment of cellular respiration measured under mitochondrial stress condition in cells repolarised as indicate. M0, M (LPS/IFNγ), M (IL-4), M (LPS/IFNγ)+(IL-4), M (IL-4)+(LPS/IFNγ) with and without HBD3.

**b:** Dot plots and relative histograms of the different patterns of expression of MHCII/CD86 and CD273/CD206 surface markers for data shown in Figure 7B. The dot plots show: 91.6%, 35.4%, 77.4%, 67.6%, 16.8%, 19.3%, 61.2, 25.6% double positive MHCII/CD86 population observed respectively in M (LPS/IFNγ), M (LPS/IFNγ+HBD3), M (LPS/IFNγ)+(IL-4), M (LPS/IFNγ)+(HBD3+IL-4), M (IL-4), M (IL-4+HBD3), M (IL-4)+(LPS/IFNγ), M (IL-4)+(HBD3+LPS/IFNγ); 26.6%, 96.5%, 52.5%, 87.4%, 93.3%, 90.2%, 61.3%, 93.2%, double positive CD273/CD206 population observed respectively in M (LPS/IFNγ), M (LPS/IFNγ+HBD3), M (LPS/IFNγ)+(IL-4), M (LPS/IFNγ)+(HBD3+IL-4), M (IL-4), M (IL-4+HBD3), M (IL-4)+(LPS/IFNγ), M (IL-4)+(HBD3+LPS/IFNγ). Debris and aggregates were eliminated from the analysis by forward- and side-scatter analysis.

**Supplementary Figure 7**

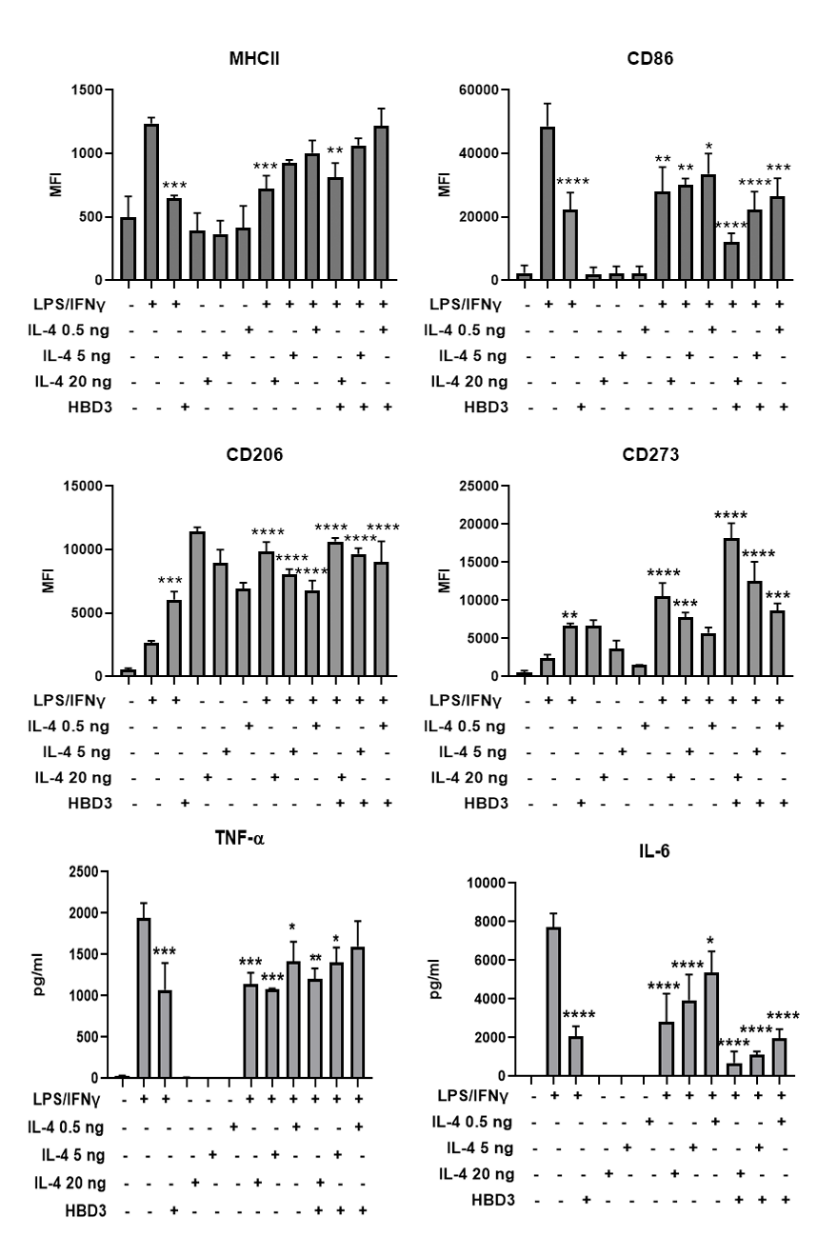

**IL-4 mimics the HBD3 effect on M(LPS+IFNγ) at 18hours**

BMDM were polarised with (LPS+IFNγ) or IL-4 20 ng, 5 ng, 0.5 ng, with or without HBD3 for 18 hours. Representative histograms of geometric mean fluorescence intensity (MFI) of MHCII, CD86, CD206 and CD273 surface markers analysed by flow cytometry. Significance values are referring to samples with LPS/IFNγ plus HBD3 +/- IL-4, in comparison to response in M(LPS+IFNγ), **** p<0.0001, *** p<0.001, ** p<0.01, * p<0.05. In the bottom, TNF-α and IL-6 expression in supernatant measured by ELISA. **** p<0.0001, *** p<0.001, ** p<0.01. Comparison were done with one-way ANOVA test.

**Supplementary Figure 8**

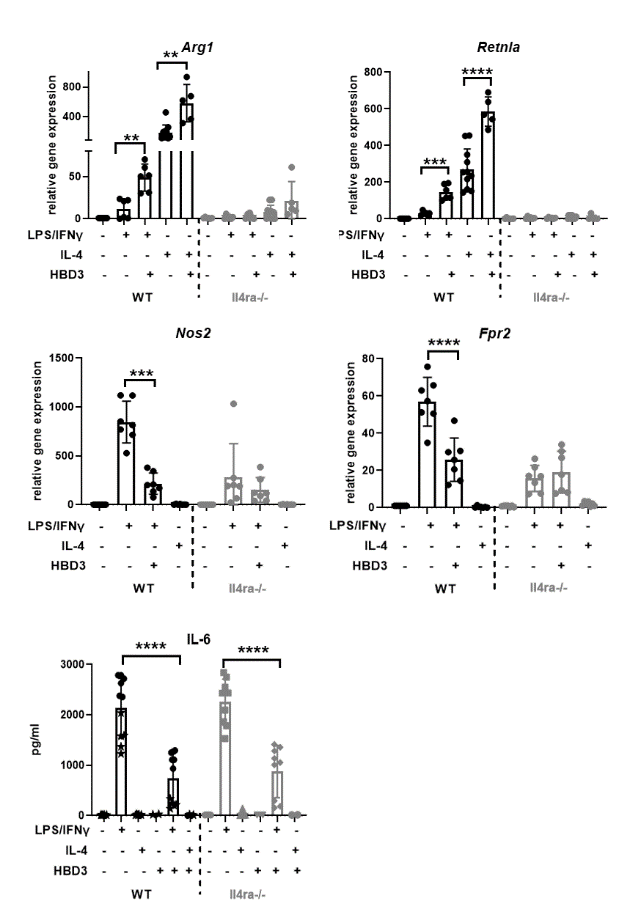

**HBD3 effect on M(LPS+IFNγ) is dependent on IL-4Rα**

BMDM were polarised with (LPS+IFNγ) or IL-4 20 ng, with or without HBD3 for 18 hours. *Arg1, Retnla, Nos2, Fpr2* gene expression were measured by real time. Results are normalized to mRNA expression of naïve macrophages. **** p<0.0001, *** p<0.001, ** p<0.01. In the bottom, IL-6 expression in supernatant was measured by ELISA. **** p<0.0001. Comparison were done with one-way ANOVA test.

**Supplementary Table 1**

| Cell | Polarised/repolarised  with | Cell name | TNFα | IL-6 | IL-10 | *Il4* | MHCII/CD86 | CD206/CD273 | *Arginase* | ATP Respiration |
| --- | --- | --- | --- | --- | --- | --- | --- | --- | --- | --- |
| **M0** | **-** | **M0** | **-** | **-** | **-** | **-** | **-** | **-** | **-** | **++++** |
| **M0** | **HBD3** | **M(HBD3)** | **-** | **-** | **-** | **-** | **-** | **-** | **-** | **++++** |
| **M0** | **LPS/IFNγ** | **M(LPS/IFNγ)** | **+++** | **++++** | **++++** | **-** | **++++** | **++** | **-** | **-** |
| **M0** | **LPS/IFNγ+HBD3)** | **M(LPS/IFNγ+HBD3)** | **+** | **++** | **+++** | **++++** | **++** | **++++** | **+** | **++** |
| **M0(*Il4ra*^-/-^)** | **LPS/IFNγ+HBD3** | **(*Il4ra*^-/-^) M(LPS/IFNγ+HBD3)** | **+++** | **++** | **ND** | **-** | **++++** | **++** | **-** | **ND** |
| **M0** | **IL-4** | **M(IL-4)** | **-** | **-** | **+** | **-** | **+** | **+**  **M(IL-4 5ng)** | **++** | **++++** |
| **M0** | **IL-4+HBD3** | **M(IL-4+HBD3)** | **-** | **-** | **++** | **+++** | **+** | **+++**  **M(IL-4 5ng)+HBD3** | **++++** | **++++** |
| **M (IL-4)** | **LPS/IFNγ** | **M(IL-4)+(LPS/IFNγ)** | **+++++** | **++++** | **ND** | **-** | **++** | **+** | **-** | **-** |
| **M (IL-4)** | **HBD3+LPS/IFNγ** | **M(IL-4)+(HBD3+LPS/IFNγ)** | **+** | **-** | **ND** | **+** | **++** | **++** |  | **++** |
| **M (LPS/IFNγ)** | **IL-4** | **M(LPS/IFNγ)+(IL-4)** | **++** | **+** | **ND** | **-** | **++** | **-** | **-** | **-** |
| **M (LPS/IFNγ)** | **HBD3+IL-4** | **M (LPS/IFNγ)+( HBD3+IL-4)** | **++** | **+** | **ND** | **+++** | **++** | **+** | **-** | **-** |

**Influence of HBD3 on macrophage responses under different polarisation conditions.**

Summary of data presented here on protein or RNA levels of cytokines, cell surface markers, Cell surface activation markers and ATP respiration during mitochondrial stress as a measure of OXPHOS following polarisation of C57Bl/6 BMDM under various conditions. ND (not done) and – expression negative or very low.
